# Dopamine and serotonin transients predict depressive symptom relief following deep brain stimulation of human subcallosal cingulate cortex

**DOI:** 10.64898/2025.11.28.691184

**Authors:** Blair R. K. Shevlin, Qi Xiu Fu, Seth R. Batten, Brian H. Kopell, Arianna Neal Davis, Matthew Heflin, Stephen Heisig, Christopher J. Rozell, Ayaka Kato, Kaustubh R. Kulkarni, Ki Sueng Choi, Ha Neul Song, Shannon O’Neill, Tanya Nauvel, Isha Trivedi, Jason Patrick White, Terry Lohrenz, Martijn Figee, Thomas Twomey, Rosalyn Moran, Dan Bang, Alec E. Hartle, W. Matthew Howe, Kenneth T. Kishida, Leonardo S. Barbosa, Vincenzo G. Fiore, Ignacio Saez, P. Read Montague, Helen S. Mayberg, Xiaosi Gu

**Author notes:** Corresponding Author: Xiaosi Gu. P. Read Montague, Ph.D., Fralin Biomedical Research Institute at VTC 2 Riverside Circle, Roanoke, VA 24016, Helen S. Mayberg, MD, Icahn School of Medicine at Mount Sinai One Gustave L. Levy Place, New York, NY 10029, Xiaosi Gu, Ph.D., Department of Psychiatry, Yale School of Medicine 40 Temple Street, New Haven, CT 06510. These authors contributed equally to this work (Blair R. K. Shevlin, Qi Xiu Fu, P. Read Montague, Helen S. Mayberg, Xiaosi Gu).

## Abstract

Recent advances in deep brain stimulation (DBS) of the subcallosal cingulate (SCC) show promise in mitigating the symptoms of treatment-resistant depression (TRD) in humans^1–3^. Monoamines, such as dopamine and serotonin, mediate the effects of pharmacological treatments of depression. However, their roles in recovery following DBS remain elusive, largely due to technical limitations of measuring these neurotransmitters in the living human brain. Here, by leveraging machine learning-enhanced electrochemistry^4–7^, we show that dopamine and serotonin signaling following DBS to the SCC predicted later depressive symptom relief in humans with TRD. We found that both dopamine and serotonin levels increased following subtherapeutic intraoperative SCC stimulation, with each neurotransmitter showing selective responses to distinct decision-making tasks. Furthermore, acute dopamine increases predicted later mood improvements during a social decision-making task, while serotonin enhancement predicted faster responses during a non-social learning task longitudinally. Critically, changes in dopamine and serotonin levels during the social decision-making task jointly predicted depressive symptom remission at 6-month follow-up. These findings illustrate the contribution of both dopamine and serotonin signaling in predicting behavioral improvement and depressive symptom remission in humans with TRD. Such neurochemical plasticity may serve as potential mechanistic biomarkers for SCC DBS mechanism and TRD treatment response.

**Significance statement:**

- Dopamine and serotonin levels increased following acute DBS to the SCC in humans.
- Acute dopamine and serotonin changes predicted later mood and response speed changes.
- Sustained TRD recovery was predicted by acute increases in both dopamine and serotonin estimates.

## Main

Deep brain stimulation (DBS) to the subcallosal cingulate (SCC) is a promising intervention for treatment-resistant depression (TRD), with robust and reproducible acute antidepressant effects observed following initial intraoperative stimulation ^2,8–10^. SCC DBS targets a critical juncture of white matter bundle tracts connecting cortical and subcortical regions, including the prefrontal cortex, hippocampus, amygdala, striatum, and the raphe nucleus ^11–16^. Related to its anatomical centrality, the SCC plays a central role in affective and cognitive functions such as emotion regulation, conflict monitoring, and reward processing ^17–20^. The SCC also exhibits distinct structural and functional alterations in depressive disorders and has thus been a key target for DBS for depression ^21–26^.

Beyond its integrative connectivity, the SCC also serves as a key neuromodulatory hub, directly influencing dopamine (DA) and serotonin (5-HT) transmission via its connections with the mesolimbic system such as ventral tegmental area and the raphe nuclei, respectively ^21,27,28^. Research on post-mortem brains has provided further support for this neuromodulatory significance, revealing associations between mood disorders and alterations in monoaminergic receptors and transporters in the SCC ^27,29,30^. These neurochemical characteristics underscore the SCC as a valid target for both clinical intervention and neurochemical investigation. The anterior cingulate expresses high densities of 5-HT transporter and receptors ^31,32^, and clinical responders to 5-HT reuptake inhibitors exhibit metabolic changes in the SCC absent in non-responders ^33,34^. Preclinical research further suggests that the antidepressant effects of chronic DBS depend on intact serotonergic pathways ^35–41^. In parallel, prior work has shown that electrical stimulation of the infralimbic cortex—the rodent homolog of the SCC—induces DA release in the striatum and nucleus accumbens ^42–47^. Collectively, these findings suggest that DBS to the SCC may modulate disruptions in monoaminergic signaling characteristic of depression ^2,48^. However, methodological constraints have limited direct neurochemical measurements in humans, forestalling a precise delineation of the roles DA and 5-HT play in the antidepressant effects of DBS treatments.

To overcome these challenges, we leveraged recent developments in machine learning-enhanced electrochemistry, a technique that allows us to measure sub-second fluctuations in extracellular monoamine levels in the brains of conscious humans as they perform cognitive tasks ^4–7,49^. Specifically, we used carbon fiber electrodes to record changes in DA and 5-HT release in the SCC along the routine insertion trajectory in humans who underwent bilateral DBS implantation for TRD (n = 10). Estimates of DA and 5-HT in the right SCC were obtained as participants performed two decision-making tasks, both before and after sub-clinical level stimulation (∼2mA) to the left SCC (**Fig. 1b**); the tasks were a non-social learning task (reversal learning) and a social exchange game (ultimatum game). The electrochemistry protocol involved repeated delivery (10 Hz) of a triangular voltage to the carbon-fiber microelectrode and measurement of induced electrochemical reactions as changes in current at the carbon-fiber tip at a sub-second scale; *in vivo* changes in DA and 5-HT levels were then estimated using a machine learning-based predictive model trained *in vitro* on large, carefully controlled wet-lab datasets (see **Methods** for details).

**Fig. 1:**
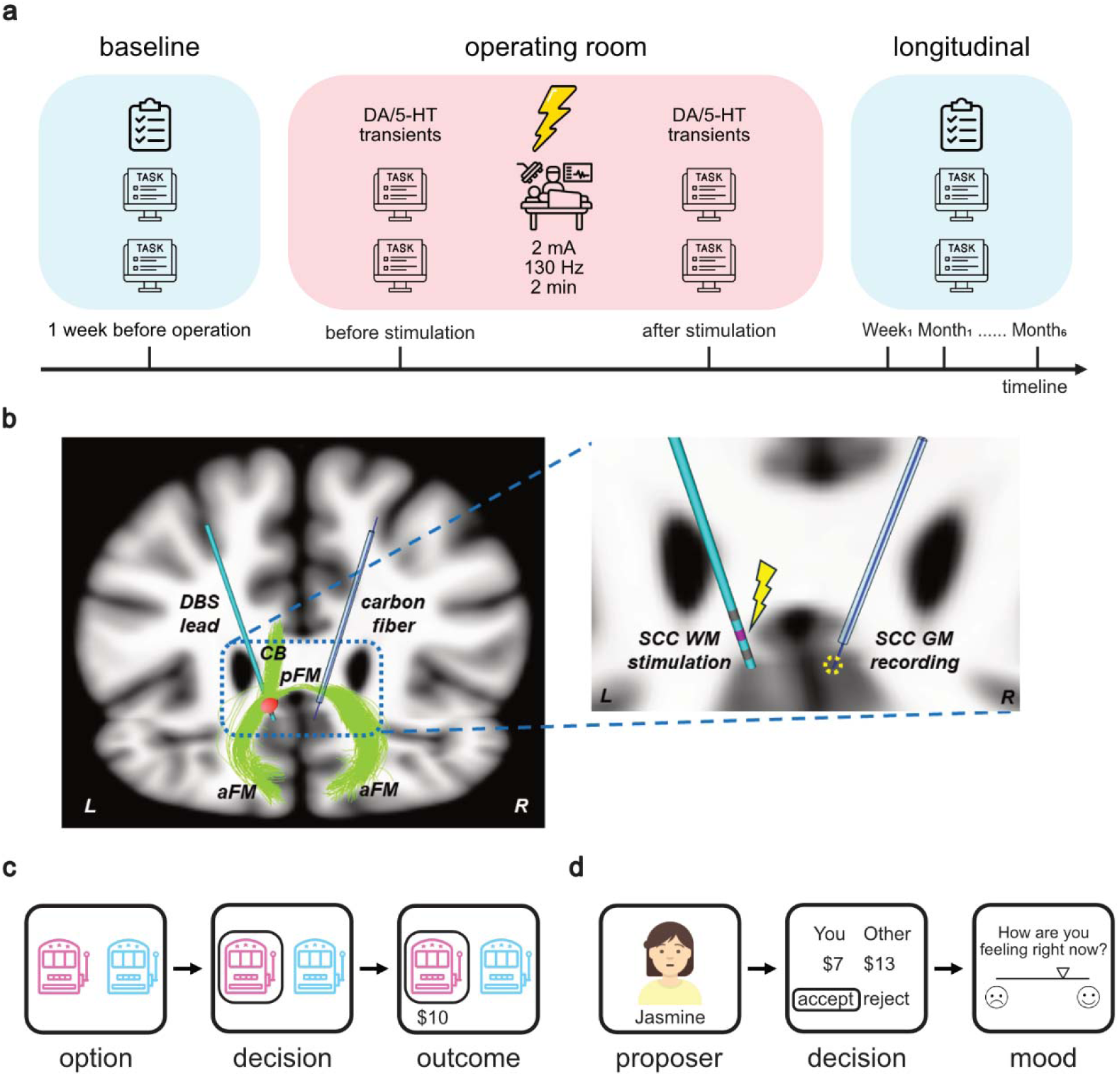
Intra-operative experimental protocol. **a,** Study design and intraoperative protocol. Overall study design included three phases: pre-operative behavioral baseline (1 week before surgery), intra-operative pre and post stimulation and recording, and post-operative assessments (1 week after surgery and monthly up to 6 months). Clinical and experimental measurements utilized for analysis included baseline HDRS, and behavioral tasks performance. **b**, Stimulation and recording protocol. Acute subtherapeutic stimulation was delivered to the left hemisphere (2 mA, 130 Hz, 90 µs, 2 min) at the subcallosal cingulate white matter (SCC WM) and intraoperative electrochemical recording was measured in the right SCC gray matter (GM). The SCC WM targets were identified using a patient-specific tractography-guided targeting algorithm. The DBS lead was implanted in the left SCC WM that stimulates the ipsilateral cingulum bundle, forceps minor and uncinate fasciculus/ventral amygdala-fugal pathway. A recording carbon fiber electrode was inserted in the lower bank of the right SCC GM for electrochemical recordings during tasks performance (-0.4∼1.3 V, 400 V/s, 10 Hz) and removed before replacing it with an electrode for bilateral DBS. The volume of tissue activated (red ball in the left hemisphere) from the left DBS lead was estimated using the following parameters: 3rd contact from the bottom, 130Hz, 90μs, and 2mA. The carbon fiber recording site (red dashed circle) is located in the right SCC GM. The forceps minor connects the two hemispheres including the two SCC regions, thus linking the left site of stimulation to the right site of recording. **c,** Reversal learning task (105 trials) with 3 blocks (35 trials/block). On each trial, participants selected one of two slot machines, and feedback was provided. From the feedback, participants could learn which machine was more likely to give the better reward across three block types: reward ($0 or $10), mixed (-$10 or $10), or punishment (-$10 or $0). The better reward outcome was probabilistic (80% for the better machine and 20% for the worse machine), with 2 reversals within each block. **d,** Ultimatum game (30 trials) with mood rating. In this task, participants needed to accept or reject an unfair split of $20 made by a virtual avatar. In 33% of the trials, participants were prompted to rate their mood from bad (0) to good (100). SCC, subcallosal cingulate cortex; DBS, deep brain stimulation; HDRS, Hamilton Depression Rating Scale. CB, cingulum bundle; aFM: anterior forceps minor; pFM: posterior forceps minor. Illustration icons are provided by Flaticon (www.flaticon.com).

We found increases in DA and 5-HT estimates in the right SCC following acute, subtherapeutic electrostimulation to the left SCC in patients with TRD. Specifically, overall DA increased during both the reversal learning and social exchange tasks, while 5-HT only increased during the social exchange task. Moreover, we found that intraoperative DA increases predicted enhanced mood during the social exchange task between one and three months after surgery, while intraoperative 5-HT increases predicted faster response times during the reversal learning task between three- and six-months post-operation. Importantly, intraoperative DA and 5-HT changes during the social exchange task jointly predicted depressive symptom relief at six-month follow-up post-surgery. Taken together, these results demonstrate how acute stimulation-induced changes in monoaminergic processes can inform our understanding of the dynamic recovery of TRD patients receiving therapeutic SCC DBS.

### Task-based changes in DA and 5-HT following acute SCC stimulation

Participants completed two repeats of the social and non-social decision-making tasks during surgery, once before acute stimulation and one once after. Specifically, participants first completed the reversal learning task and then the ultimatum game.

Next, each participant received two minutes of subtherapeutic, unilateral electrostimulation (2 mA, 90 µsec, 130 Hz, single-blind) to the left SCC. After stimulation, participants again performed the ultimatum game, followed by the reversal learning task.

During the reversal learning task (**Fig. 1c**), participants completed 35 trials in each of three blocks with distinct outcomes (total trial *n* = 105): reward blocks with $10 and $0 outcomes, punishment blocks with $0 and -$10 outcomes, and mixed blocks with $10 and -$10 outcomes. In each trial, participants selected one of two machines that gave probabilistic outcomes of $10, $0, or -$10. The better machine was always associated with 80% winning probability, while the less rewarding machine was rewarded with 20% probability. The machine with the higher winning probability switched every 12 or 13 trials. The timing of the reversals was pseudorandomized across participants.

During the ultimatum game (**Fig. 1d**), participants completed 30 trials with a series of one-shot interactions with simulated human partners. In each trial, participants were asked to accept or reject an unfair split of $20 (ranging from $1 to $9). If they accepted the offer, both the participant and the simulated proposer would receive the proposed amounts. If they rejected the offer, both parties would receive nothing. All participants received the same offers, but in a randomized order. On a third of trials, participants were also asked to rate their current mood by moving a slider along a visual analog scale ranging from negative (sad emoji) to positive (happy emoji).

Estimated neurotransmitter levels were defined as the sum of samples within a 500 ms window following outcome reveal in the reinforcement learning task and offer presentation in the ultimatum game (**Extended Data Figure 1**). To characterize changes in DA and 5-HT across pre- and post-stimulation periods, we ran mixed-effect general linear models (GLMs) for each task and each neurotransmitter (see Methods for details) in which we predicted participant-level estimates of overall DA or 5-HT levels using session (pre-stimulation = 0, post-stimulation = 1), normalized trial number (z-scored within task), and their interaction.

For the reversal learning task (**Fig. 2a**), DA estimates following outcome reveal significantly increased after stimulation (β = 6.41, *t* = 2.37, *P* = 0.045), while 5-HT estimates did not (β = 1.31, *t* = 0.37, *P* = 0.722). At the subject level, we observed increases in overall DA following stimulation among six (out of eight) participants, but only four participants showed post-stimulation increases in 5-HT (**Fig. 2c**; Supplementary Table 3). DA and 5-HT estimates during the reversal learning task did not significantly vary with block type (*P*s > 0.05, **Fig. 2e**) or other task features (Supplementary Table 6-9).

**Fig. 2:**
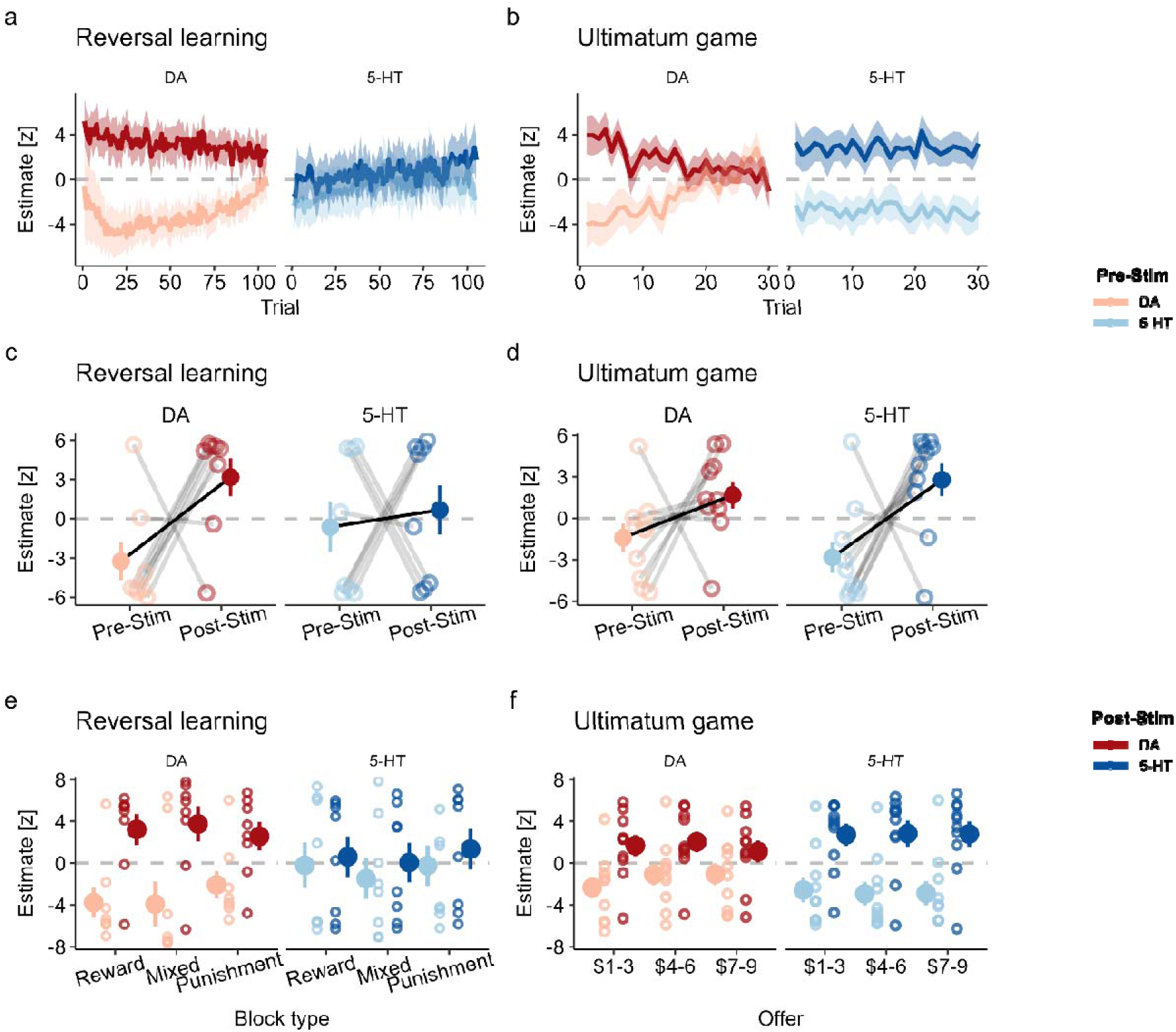
Stimulation-induced changes in neurotransmitters are task-dependent. **a,** Stimulation-induced changes in neurotransmitter estimates during the reversal learning (RL) task. Dopamine (DA) estimates increased significantly following stimulation (mixed-effects linear regression: β = 6.41, t = 2.37, P = 0.045), whereas serotonin (5-HT) estimates did not change (β = 1.31, t = 0.37, P = 0.722). Shaded regions represent SEM across participants. **b,** Stimulation-induced changes in neurotransmitter estimates during the ultimatum game (UG). Serotonin (5-HT) estimates increased significantly following stimulation (β = 5.62, t = 2.57, P = 0.028), whereas dopamine (DA) estimates did not change (β = 3.09, t = 1.66, P = 0.129). **c,** Individual participant responses during RL. Six of eight participants showed increased DA following stimulation, while four of eight showed increased 5-HT. Open circles represent individual participant means; filled circles represent group means. Error bars represent SEM across participants. Gray lines connect pre- and post-stimulation values for each participant. **d,** Individual participant responses during UG. Eight of ten participants showed increased DA following stimulation, and eight of ten showed increased 5-HT. **e,** Neither DA nor 5-HT estimates were modulated by RL task features (reward, mixed, or punishment blocks) before or after stimulation (all Ps > 0.05). **f,** Neither DA nor 5-HT estimates were modulated by UG offer amounts ($1-3, $4-6, or $7-9) before or after stimulation (all Ps > 0.05).

For the ultimatum game, 5-HT estimates during the offer period significantly increased after stimulation (β = 5.62, *t* = 2.57, *P* = 0.028), while DA estimates during this period did not (β = 3.09, *t* = 1.66, *P* = 0.129; **Fig. 2b**). Here, eight (out of ten) participants showed increases in either DA or 5-HT following stimulation (**Fig. 2d**; Supplementary Table 4). Neither DA nor 5-HT estimates were significantly related to offer amount (*P*s > 0.05, **Fig. 2f**) or other task features (Supplementary Table 10-20).

These task-dependent neurotransmitter changes were further supported by directly testing for a task-by-stimulation interaction in an additional mixed-effect GLM (see Supplementary Table 5).This analysis demonstrated that post-stimulation changes in DA during the reversal learning task were greater than during the ultimatum game and that 5-HT changes during the ultimatum game were greater than during the reversal learning task (DA task × interaction: β = -2.89, *t* = -9.28, *P* < 0.001; 5-HT task × interaction: β = 4.49, *t* = 15.90, *P* < 0.001). Finally, we observed no significant stimulation-induced changes in estimated norepinephrine levels during either task (see **Extended Data Fig. 2**). Together, these results suggest that the acute neurochemical effects of SCC DBS may be selective for DA and 5-HT, depending on the task context.

### Neurochemical responses to stimulation track longitudinal behavioral changes

Next, we sought to examine how these acute changes in DA and 5-HT might relate to task-based behavioral changes in the participants. Therapeutic stimulation was turned one day after surgery for all participants, and therapeutic stimulation was administered over six months. In addition to the surgical session, the two behavioral tasks were administered during a pre-operative baseline session, a one-week post-operative session, and monthly follow-up visits for six months. We calculated task-based behavioral changes by contrasting each follow-up session with baseline performance during the pre-operative session. For the reversal learning task, this included average optimal choice frequency (i.e., choosing the option with a higher win probability) and mean log-transformed response times (log RT). For the ultimatum game, this included average acceptance rates, mean log RT, and mood ratings.

Over the six-month treatment period, we observed that RTs, a measure of psychomotor speed, decreased for both tasks (**Fig. 3a, c**). These faster response speeds were observed as early as Month 1 (paired-samples *t-*test, reversal learning: *P* = 0.002; ultimatum game: *P* = 0.008) and were sustained through Month 6 (reward learning: *P* = 0.005; ultimatum game: *P* = 0.011). While there were no significant changes in choice behavior during the reversal learning task (*P*s > 0.05; **Fig. 3b**), participants were less likely to accept unfair offers during the ultimatum game (*P* = 0.015; **Fig. 3d**) and more likely to report an elevated mood during the social exchange task over time (*P* = 0.022; **Fig. 3e**). Notably, these latter patterns were only statistically significant during the sixth month of their DBS treatment.

**Fig. 3:**
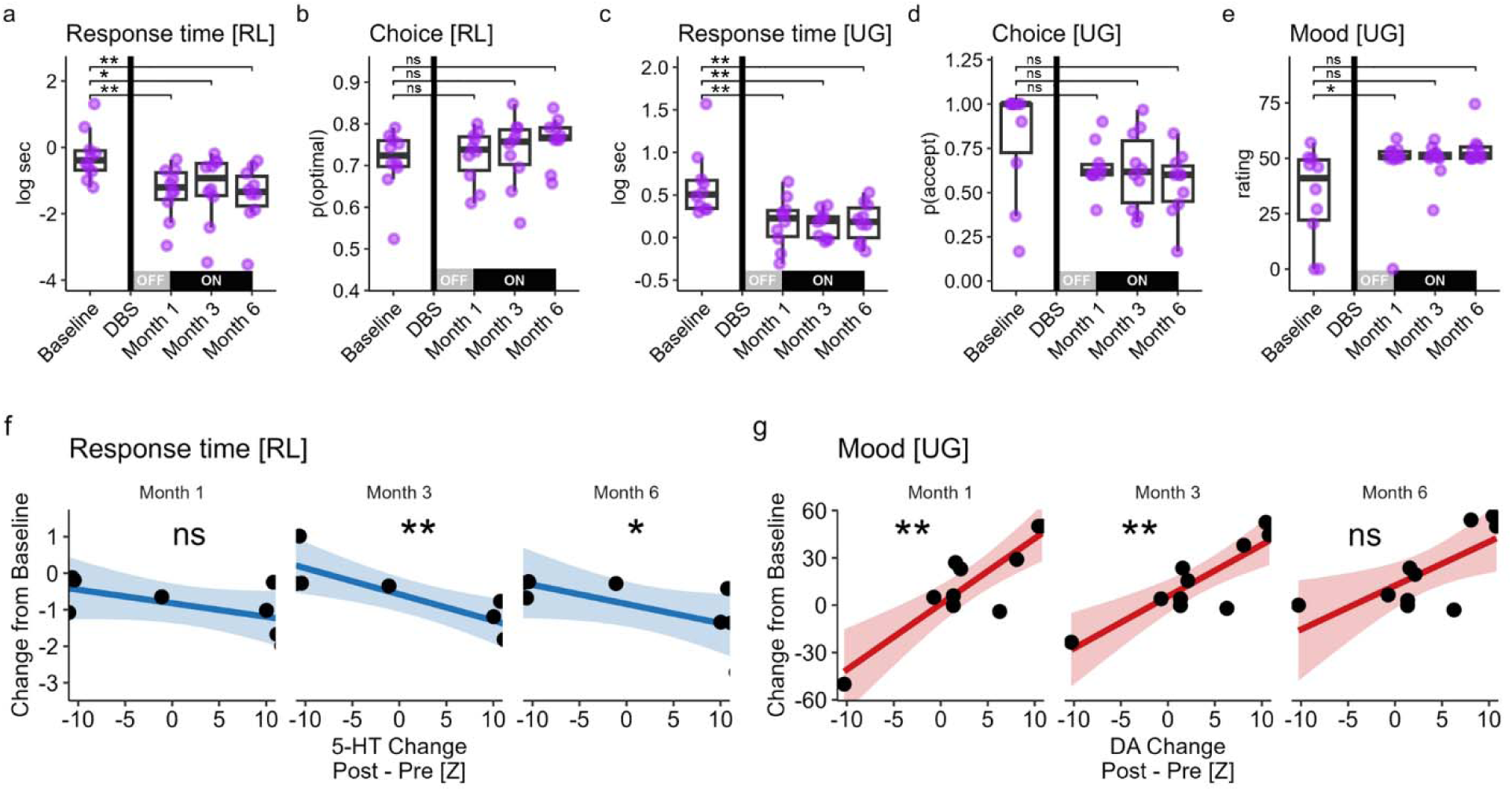
Neurotransmitter correlates with behavioral changes following DBS treatment. **a,** Longitudinal trends in response times (RT) during the reversal learning task (RL). Compared to baseline, participants’ RT was significantly reduced in Month 1 (Paired-samples *t = -*4.24, *P* = 0.002), Month 3 (*t* = -2.77, *P* = 0.022), and Month 6 (*t* = -3.64, *P* = 0.005). **b,** Optimal choice patterns during the RL task showed no significant longitudinal trends. **c,** Longitudinal trends in RT during the social exchange task (UG) showed significant reductions compared to baseline (Month 1: *t* = -3.36, *P* = 0.008; Month 3: *t* = -3.93, *P* = 0.003; Month 6: *t* = -3.18, *P* = 0.011). **d,** Following chronic DBS, selfish behavior increased and participants were less likely to accept offers in Month 6 compared to baseline (*t* = -3.00, *P* = 0.015). **e,** Compared to baseline, participants’ mood ratings increased in Month 6 (*t* = 2.76, *P* = 0.022). **f,** Correlations (Spearman’s rho) between stimulation-induced changes in 5-HT during RL and longitudinal shifts in RT. RT change is the difference in log-transformed RT from baseline. By Month 3, larger increases in post-stimulation 5-HT predicted faster RT (rho = -0.88, *P* = 0.007), a pattern which persisted in Month 6 (rho = -0.76, *P* = 0.037). **g,** Correlations between stimulation-induced changes in DA during UG and longitudinal shifts in mood ratings. Mood change is the difference in average mood ratings from baseline. Starting in Month 1, larger increases in post-stimulation DA predicted larger shifts in self-reported mood (rho = 0.68, *P* = 0.030). This pattern was also significant in Month 3 (rho = 0.67, *P* = 0.035) but not Month 6 (rho = 0.50, *P* = 0.143). Asterisks represent statistically significant changes (*^ns^P* > 0.05, \**P* < 0.05, \*\**P* < 0.01).

We then correlated these longitudinal changes in task behaviors with stimulation-induced changes in neurotransmitter estimates (post-stimulation minus pre-stimulation). For the reversal learning task, we found that greater post-stimulation increases in 5-HT levels were significantly correlated with faster RT starting in Month 3 (**Fig. 3f**; Spearman’s rho: -0.88, *P* = 0.007) and continuing through Month 6 (Spearman’s rho: - 0.76, *P* = 0.037). For the ultimatum game, we found that greater post-stimulation increases in DA levels were significantly correlated with greater increases in mood ratings starting in Month 1 (**Fig. 3g**; Spearman’s rho: 0.79, *P* = 0.007), and continuing in Month 3 (Spearman’s rho: 0.77, *P* = 0.009) and marginally in Month 6 (Spearman’s rho: 0.64, *P* = 0.054). Together, these results suggest that post-stimulation 5-HT (but not DA) changes were primarily related to changes in response speed longitudinally, whereas post-stimulation DA (but not 5-HT) changes predicted later mood improvements in these TRD patients.

### Sustained recovery predicted by increased DA and 5-HT estimates

Finally, to examine the potential utility of intra-operative electrochemical measures in predicting clinical outcomes, we analyzed the relationship between stimulation-induced neurochemical changes and patients’ long-term depressive symptom changes as measured by the Hamilton Depression Rating Scale (HDRS). The HDRS is a physician-administered survey and the primary outcome measure of DBS treatment response.

Baseline HDRS scores were obtained weekly during the month prior to surgery and subsequent scores were obtained during a one-week post-surgical session and monthly follow-up visits for six months. After six months of chronic DBS treatment (**Fig. 4a**), participants exhibited an average 60.9% reduction in HDRS symptom severity scores (min = 23.4%, max = 90.7%). Consistent with our previous SCC DBS trials using pre-operative tractography defined targeting for chronic therapeutic stimulation ^1,9,10,50^, seven out of ten participants met criteria for antidepressant response (i.e., greater than 50% decrease in HDRS), and five out of ten were in clinical remission (i.e., HDRS < 8; **Extended Data Table 2**). These results demonstrate that the effectiveness of the current DBS protocol was consistent with previously published trials using identical targeting methods ^10^.

**Fig. 4:**
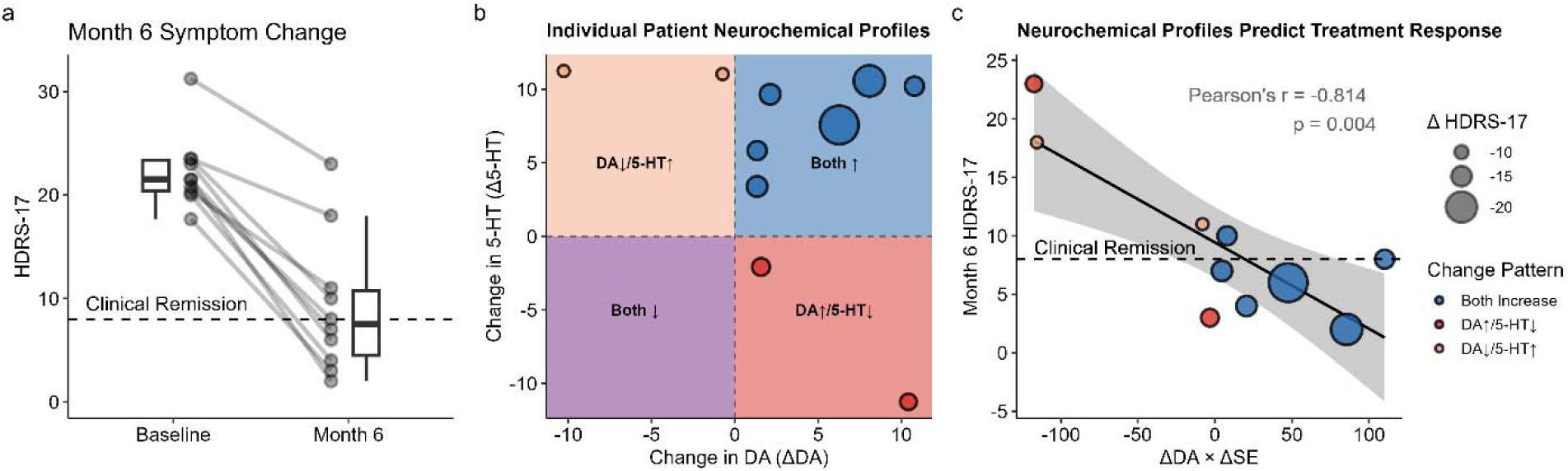
Neurotransmitter correlates of symptom improvement. **a,** Change in HDRS-17 after six months. Each individual (dot) experienced a decline in symptom severity scores following DBS treatment. **b,** Change in month 6 HDRS-17 as predicted by stimulation-induced changes in 5-HT and DA during the ultimatum game. Larger induced increases in DA paired with post-stimulation increases in 5-HT predicted reduced symptom severity (linear regression; ΔDA_Pre-Post_ x Δ5-HT_Pre-Post_: f3 = -0.14, *P* = 0.003). **c,** Strength of post-stimulation increases in 5-HT and DA were correlated with month 6 HDRS-17 scores.

Next, using the average changes in monoamine levels, and their interactions, as regressors, we sought to estimate HDRS scores at Month 1, Month 3, and Month 6. We found that a model that consisted of DA and 5-HT changes during the ultimatum game outperformed alternative models in predicting HDRS scores (Supplementary Table 21). In this model, the interaction between DA changes and 5-HT changes measured during the ultimatum game predicted lower HDRS scores during Month 1 (linear regression, *b* = -0.13, *t* = -2.68, *P* = 0.037) and Month 6 (*b* = -0.14, *t* = -4.86, *P* = 0.003). Specifically, HDRS scores improved the most when DA and 5-HT both increased after SCC stimulation (**Fig. 4b**). We did not identify significant relationships between neurochemical changes measured during the reversal learning task and HDRS scores at any time point (Month 1: *b* = -1.00, *t* = -2.52, *P* = 0.088; Month 3: *b* = -1.04, *t* = -2.43, *P* = 0.072; Month 6: *b* = -0.64, *t* = -0.91, *P* = 0.416). Comparisons of alternative model specifications are provided in Supplementary Table 21. Together, these results suggest that successful responses to DBS treatment were predicated upon joint changes in DA and 5-HT estimates specifically during the social exchange task (**Fig. 4c**).

## Discussion

Deep brain stimulation to the SCC has widespread neurobiological and clinical effects ^1,10,11,50,51^. Here, we leveraged machine learning-enhanced electrochemistry to measure, for the first time, sub-second fluctuations in neurotransmitter dynamics following acute electrostimulation to the SCC in awake humans with TRD. We observed functional relationships between acute, stimulation-induced changes in monoamines release and longitudinal changes in both task behaviors and depression symptom severity. Specifically, we found increased DA levels during a reversal learning task and increased 5-HT levels during a social exchange task (ultimatum game) following subtherapeutic intraoperative stimulation. Acute DA increases were associated with later improvement in mood during the ultimatum game, while acute 5-HT increases were associated with faster response speeds during the reversal learning task longitudinal. Critically, DA and 5-HT responses to acute stimulation jointly predicted longitudinal DBS treatment success. Together, these results provide the first intracranial evidence from humans supporting the critical roles of DA and 5-HT in the antidepressant response of SCC DBS.

Our observation of dopaminergic and serotonergic changes in the SCC is consistent with its known neuroanatomical profile. The neurochemical dynamics of the SCC are closely linked to a broader limbic-cortical network, which can be activated through DBS via the intersection of four key white matter bundles: the forceps minor, uncinate fasciculus, cingulum bundle, and midline frontostriatal fibers ^52,53^. These tracts facilitate dopaminergic inputs from the ventral tegmental area (VTA) to the SCC while also sending efferent projections that modulate DA release in the VTA ^54,55^. Efferents from the SCC project onto GABAergic interneurons in the dorsal raphe nucleus, modulating serotonergic projections across the brain ^56–58^. Optogenetic stimulation of infralimbic targets in the dorsal raphe nucleus drive motivated escape behaviors in rodents ^59,60^, a potential parallel to the relationship between acute changes in 5-HT changes and reductions in response times observed here. The anterior cingulate expresses high densities of 5-HT transporters ^31,32^, and reduced 5-HT 1B receptor binding potential in this region has been linked to depressive disorders ^61,62^. Similarly, targeted stimulation of the rodent analog of the SCC alters DA release in the VTA and nucleus accumbens ^46,47^ (but see ^63^) and influences 5-HT release in the hippocampus ^37,50^, highlighting the SCC’s role in regulating monoaminergic transmission ^41,48^. Complementing prior work establishing how DBS can modulate cingulate connectivity ^1,10,50^, our findings provide initial evidence that DBS also alters monoamine flows traveling along these pathways in humans. However, future work using non-human animal models is needed to illustrate the exact pathways and mechanisms through which neurotransmitters in the SCC are changed by stimulation.

Our findings also align with established roles of DA and 5-HT in reward learning ^64,65^ and social behaviors ^66–68^, respectively. However, most prior research has been based on animal models or indirect inferences from human neuroimaging or pharmacological studies. More recently, a handful of human studies using a similar neurochemistry approach have demonstrated that DA and 5-HT dynamics in the basal ganglia track various task-dependent rewards ^4–7,69,70^ and sensory signals ^49,71^ in patients undergoing DBS for movement disorders. Importantly, in vivo validation for this method has been provided using optogenetics in rodents using the same carbon fiber electrodes used in humans ^6^. In contrast to similar experiments conducted with human participants ^4,4,5,7,69,70^, we did not find a direct relationship between estimated DA or 5-HT levels (e.g., z-scored values summed within a 500 ms window following task-relevant events) and trial-level value and learning signals either before or after stimulation (Supplementary Tables 6-9, 10-20). We speculate that several factors may have accounted for this discrepancy, including differences in recording sites (cortical vs. subcortical) and patient populations (TRD vs. Parkinson’s disease). While both regions contribute to learning and cognition, dopamine in prefrontal circuits supports cognitive control functions such as working memory gating and maintenance ^72,73^, whereas striatal dopamine supports the acquisition and updating of action values through reinforcement learning ^64,74^.

Similarly, serotonin has been suggested to mediate neural plasticity in the prefrontal cortex and subserve executive functions such as working memory and behavioral flexibility ^75,76^, which is distinct from its role in subcortical regions ^77,78^. Finally, although direct comparisons are scarce, TRD and Parkinson’s are distinct conditions characterized by separate neurobiological mechanisms; as such, the neurochemical milieu of individual brains affected by these conditions are drastically different, which may have additionally contributed to the divergent findings in the current study compared to previous work.

Although the neurochemical changes did not correlate with trial-level task features or behaviors measured in the operating room, we identified multiple relationships between intraoperative DA and 5-HT changes and task-based behavioral changes measured longitudinally. Specifically, we found that post-stimulation DA increases during the ultimatum game were associated with later improvement in mood ratings at months one and three, while post-stimulation 5-HT increases during the reversal learning task were linked to faster reaction times during the same task at months three and six. This lag in behavioral relevance of the observed neurochemical changes likely reflects delayed emergence of behavioral improvements themselves: task-based behaviors showed no acute changes following acute, intraoperative stimulation (Supplementary Figure 3) but emerged gradually starting at month one, after therapeutic stimulation was initiated.

Two factors may explain this delay. First, the intraoperative stimulation amplitude (2 mA) was substantially lower than the therapeutic levels used during chronic stimulation, potentially insufficient to produce immediate behavioral effects. Future dose-response studies could clarify whether higher acute stimulation levels induce more rapid behavioral changes. Second, these decision-making tasks likely engage multiple neural circuits and substrates beyond monoaminergic signaling alone, which may require weeks to months of stimulation to facilitate behavioral adaptation. Future studies using multimodal neuroimaging and denser behavioral sampling could directly test this hypothesis. Overall, the identified brain-behavior relationships are consistent with prior research linking dopaminergic dysfunction to depressive symptoms related to mood and motivation ^54,79,80^ and demonstrate that interventions targeting serotonergic pathways alleviate psychomotor deficits in depression ^81,82^.

The clinical outcomes we observed align with previous tractography-guided SCC DBS studies ^1,3,52^. Importantly, the current study extends our understanding of recovery by linking clinical improvement to acute changes in both dopaminergic and serotoninergic systems, with potential for identification of TRD phenotypes relevant to treatment optimization broadly. Notably, greater stimulation-evoked changes in DA and 5-HT levels during the ultimatum game predicted lower HDRS scores at both early (month one) and later (month six) phases of SCC DBS treatment. These results suggest that successful, sustained DBS responses may depend on simultaneous changes in both DA and 5-HT systems, highlighting the importance of individual differences in monoaminergic plasticity for treatment outcomes ^83^. Related work has consistently found that responders to DBS treatment show normalization in the structure and function of limbic-cortical depression brain networks ^50,84,85^ – changes which may reflect transsynaptic alterations or activity dependent plasticity effects in the stimulated regions ^86,87^. Future work should examine whether identified DA and 5-HT changes might map onto network-level differences linked to the SCC in individuals with TRD. Together, the current findings underscore the role of monoamines in TRD recovery and highlight the potential for leveraging neurochemical metrics for treatment optimization.

The current findings should be interpretated with the following limitations in mind. First, the TRD sample studied here is small and homogenous in terms of their symptom severity; further work is needed to clarify the generalizability of these findings in larger patient cohorts with a broader range of symptoms. A larger, more heterogenous sample would also allow us to further cross-validate the identified relationships between neurochemical plasticity and clinical improvement. Second, due to constraints of the surgical procedure, the current study only examined the effects of left-sided stimulation on contralateral neurochemical dynamics; future work should also investigate laterality effects, which has been implicated in depression pathophysiology and DBS efficacy ^88–92^. These surgical constraints, combined with the small sample size, also limited our ability to fully balance task order in the current study. Third, although voltametric data were collected during stimulation, the neurochemical signals (nanoampere-level) were severely degraded by electrical interference from concurrent stimulation (milliampere-level), rendering them unanalyzable. Consequently, we could only compare neurochemical measurements obtained before versus after - but not during - intraoperative stimulation. This limitation leaves unresolved whether the observed monoamine changes were triggered by stimulation onset or occurred only after stimulation offset. Determining the causal relationship between stimulation and DA/5-HT dynamics will require future electrochemical advances that enable artifact-free concurrent stimulation and recording. Despite these constraints, supplementary analyses ruled out learning or repetition effects as explanations for the observed neurotransmitter changes, supporting their interpretation as stimulation-specific, task-dependent responses (Supplementary Figure 2). Finally, the DA and 5-HT estimates were inferred by machine learning and were indirect by nature. Extensive in vitro and in vivo validation, including successful separation of mixture data^5,6,49^, demonstrates the method’s capacity to monitor neurochemicals in the living human brain. Nonetheless, the unknown nature of each individual brain’s neurochemical milieu in vivo still presents challenges that require future technical advances in human neuroscience methods.

In summary, our findings offer new insights into the role of DA and 5-HT within the limbic-cortical depression network targeted by SCC DBS. The observed neurochemical changes represent a type of neurochemical plasticity that is potentially crucial for both advancing our understanding of DBS mechanisms and treatment optimization in the future.

## Method

### Participants and clinical assessments

Ten consecutive participants with treatment-resistant major depressive disorder (TRD) were recruited to participate in this study via their enrollment in one of two single-site open-label SCC DBS clinical trial studies using two prototype DBS devices under physician sponsored FDA IDE G130107. Five were implanted with a Medtronic Summit RC+S device (ClinicalTrials.gov identifier <u>NCT04106466);</u> five with a Medtronic Percept PC device (<u>NCT05773755)</u>. The DBS protocol (IRB protocol number for RC+S device: #19-01002; for PC device: #22-01731) and the intraoperative electrochemical data collection procedure (IRB #13-00415) were approved by the Institutional Review Board (IRB) at the Icahn School of Medicine at Mount Sinai. The data analysis protocol of the deidentified electrochemical data collection was approved by the Virginia Tech IRB (#11-078). All participants identified as non-Hispanic White with an even split in sex (5 females, mean ± SE = 44.0 ± 4.20 years). A summary of participants’ characteristics is provided in **Extended Data Table 1**. Participants consented to the two data collection protocols independently and their ability of enrolling in one study did not dependent on the other. For the electrochemistry protocol, participants were informed that the study involved a research-exclusive probe (carbon-fiber electrode) and an extra time (maximum 60 minutes) during their surgical implantation. Participants did not receive compensation for study participation or for earning points for the behavioral tasks. No adverse or unanticipated events occurred during or as a result of the intraoperative electrochemical experiments or related behavioral experiments.

Clinical symptoms severity was assessed by an independent rater using the 17-item HDRS, MADRS and self-reported Beck Depression Inventory during weekly visits to the study site, among other behavioral scales and testing. Participants met weekly with the study psychiatrist for evaluation of symptoms and stimulation adjustments if needed using a combination of HDRS-17 scores relative to the previous week and their clinical judgement. According to established study criteria, a decrease in HDRS-17 scores greater than 50% of the presurgical average was set as the threshold for “response” (4 responders in RC+S cohort and 3 responders in PC cohort (see **Extended Data Table 2**). Remission was defined as HDRS-17 < 8 and MADRS < 10. Baseline HDRS-17 scores reflect the average of four weekly ratings obtained during the month prior to surgery. We reported the analysis of electrochemistry from ten participants during their acute subtherapeutic DBS stimulation (8 from RL, 10 from UG). Two participants (responders: participants 1 and 3) were excluded from analysis of the RL task due to challenges in aligning their task event and electrochemistry timelines.

### SCC DBS protocol

Bilateral electrode array leads (3387, Medtronic Summit RC+S in participants 1-5; B33015-42, Medtronic Percept PC in participants 6-10) were eventually implanted in each participant, one in left and one in right SCC, as determined from tractography described in previous study^1^. We used a patient-specific connectome-based targeting approach to identify DBS targets at the intersection of four white matter bundles: forceps minor, cingulum bundle, uncinate fasciculus, and frontostriatal fibers ^52,93^.

For the intraoperative experiment, only one DBS electrode was implanted to the left SCC. For the right hemisphere, the carbon fiber electrode was inserted along the planned DBS electrode trajectory, before the DBS electrode was implanted in that hemisphere. After the experiment, the carbon fiber electrode was removed and a second DBS electrode was placed into the same trajectory on the right side. During the intraoperative experiment, subtherapeutic acute stimulation (2mA) was delivered using current-controlled pulse generators, targeting the left cingulum bundle to engage forceps minor and uncinate fibers which provide connections to the right SCC (**Fig. 1b**). The approximate MNI coordinates for both stimulation and recording sites are provided in Supplementary Table 22.

Therapeutic DBS started the day after surgery as per the ongoing clinical trial protocols in all patients. DBS therapy consisted of bilateral monopolar high-frequency (130 Hz for RC+S cohort; 125 Hz for PC cohort) stimulation on a single contact per hemisphere with 90 µs pulse width. Patients started their therapeutic stimulation with 4.5 mA. During the 6-month observation phase, location, pulse width, and stimulation frequency remained unchanged. Dose was increased as needed from 4.5 mA to between 6mA and 8 mA in 7 of 10 patients with stable dose maintained by 4 months in 5 patients. There were no significant differences on HDRS-17 scores at baseline and across 6 months between the two cohorts (See **Extended Data Fig. 3**). The initial stimulation amplitude and at the end of the 6-month study period in each participant are listed in **Extended Data Table 2**. Note, these 10 patients are a subset of two larger TRD cohorts participating in separate SCC DBS experimental studies whose primary hypotheses do not utilize these chemical measurements; those results will be published separately.

### Study procedure

The general study design included three distinct phases (pre-operative, intra-operative, post-operative) that contained varies levels of data collection including clinical, behavioral, and electrochemical measures (**Fig. 1a**). Pre-operative baseline behavioral tasks were collected 1 week before surgery.

During the intraoperative experiment, each participant performed the behavioral tasks twice – once before and once after stimulation. A DBS electrode was first implanted targeting SCC white matter on the left hemisphere and the carbon fiber research electrode was then inserted along the planned DBS trajectory targeting the SCC grey matter on the right hemisphere for electrochemical acquisition (**Fig. 1b**). Propofol anesthesia was administered for lead implantation and discontinued at least 60 minutes before testing to allow sufficient time for patients to wake up for the experiment. After both electrodes were in position, participants also engaged in verbal communications with the research team during the pre-task preparation period. These steps ensured that participants were fully awake from anesthesia and could perform the tasks, and that anesthesia wearing off effects would not affect the recorded signals, which was supported by our analyses on the general temporal patterns of DA and 5-HT (Supplementary Figure 1). Once the intraoperative experiment began, participants went through task instructions again and completed practice trials before beginning the actual tasks as additional steps to ensure their full wakefulness and ability to complete tasks. Throughout the experiment, the participants were closely monitored by the anesthesiologist, neurosurgeon, and the research team to ensure their sustained capacity for research participation.

Participants first completed the reversal learning task followed by the ultimatum game. Next, a single-blind high-frequency stimulation was delivered to the left hemisphere using a square wave pulse at 2 mA, 130 Hz, 90 µs for 2 minutes at the predefined electrode contact to be used for chronic stimulation in each participant. After acute stimulation, participants completed the ultimatum game again, followed by the reversal learning task. Electrochemistry was recorded concurrently throughout the protocol in the grey matter of the SCC in the right hemisphere. While the voltametric current was recorded during stimulation, the magnitude of current change for the voltammetry signal was at the nA level, whereas the stimulation was at the mA level and essentially overrode the voltammetry signal. As such, the neurochemical signals during stimulation were currently unanalyzable and we focused on pre- and post-stimulation recordings.

During testing, the participant laid in a semi-upright position and viewed a computer monitor at around 100 cm (about 3.28 ft). They used a gamepad to submit their responses in both behavioral tasks. Each behavioral task took an average of 10 minutes. After completing the session, the carbon fiber electrode was removed, and the participant was re-anesthetized to complete the DBS clinical procedure (right DBS lead and pulse generator implantation). The behavioral data collection session lasted approximately 50 minutes. There were no adverse effects of these procedures.

As per the two clinical trials protocols, active bilateral therapeutic DBS was initiated the day after surgery using common stimulation parameters at the tractography defined locations in each hemisphere (130 Hz, 90 ms, 4.5 mA). Clinical assessments and standardized rating scales were performed weekly; behavioral task sessions were performed at week 1 and then monthly for 6 months (**Fig. 1a**). Behavioral tasks were collected in a laboratory space with stimulation turned off, and participants resumed their clinical stimulation after their post-operative sessions. Notably, short periods of SCC DBS discontinuation do not result in any overt behavioral effects as determined by previous discontinuation experiments ^3,94^. Summary statistics of both intraoperative and longitudinal session-level behavior across tasks are provided in Supplementary Table 2.

### Reversal learning (RL)

We used a two-arm bandit reversal learning task to study reward decision-making (**Fig. 1c**). Over three blocks, participants completed a total of 105 trials (35 trial/block). On each trial, participants selected between two slot machines with probabilistic rewards. Every trial, one machine always yielded a better outcome with an 80% probability while the other machine yielded a better outcome with a 20% probability. After selecting a machine, participants viewed a three-second slot machine animation before receiving outcome feedback for an additional 1.5 seconds. Potential outcomes were based on the block type. During practice, participants were informed about the potential outcomes. In the reward block, the outcomes were $10 or $0. In the mixed block, the outcomes were $10 or -$10. In the punishment block, the outcomes were $0 or -$10. Each block contained two reversals which were predefined and pseudorandomized at the participant-level. Participants were instructed to earn as much money as possible, and the task was self-paced.

### Ultimatum game (UG)

We used a modified, multi-trial one-shot ultimatum game to study social decision-making (**Fig. 1d**). During each session, participants completed 30 self-paced trials where a virtual avatar proposed a monetary split of $20. Participants decide whether to accept or reject the proposed monetary split. If the participant accepted the offer, they received feedback indicating that they received a portion of the proposed split. If the participant rejected the offer, they received feedback indicating that they received $0. In both cases, feedback lasted for 1 second. Offers ranged from $1 to $9 based on an experimenter-defined offer list. Offers always start at $5 followed by randomized offer amounts. Every 3 or 4 trials (i.e., a third of all trials), participants were asked to report their current mood on a Likert-scale from bad (0) to good (100).

### Electrochemistry

We applied machine learning-enhanced electrochemistry methods to participants as detailed in previous work ^4,5,69–71^. Briefly, a standard triangle voltage waveform was applied to a custom-made carbon fiber electrode inserted in the patient, and the resulting current signal was recorded (pClamp, Axon Instruments, Molecular Devices). The carbon fiber electrodes were constructed from Alpha Omega NeuroProbe Sonus guide tubes (STR-901080-10), measuring 22.7 cm in length and approximately 1.27 mm in diameter. Each electrode consisted of a glass capillary (Molex #1068450450; 25.5 cm in length, 440 µm in diameter) housing a carbon fiber measurement surface with a 7 µm diameter and a 120–180 µm in length ^5^. Detailed schematics and dimensions of the human carbon fiber electrodes are provided in Supplementary Figure 4. Multi-variate neurochemical concentration estimates (DA, 5-HT, NA, and pH) using these current traces were made using an ensemble of neural network models trained on currents from in vitro solutions containing known analyte concentrations (see **Extended Data Fig. 4**). For details on how these methods were validated in rodent models using optogenetically-expressed dopamine, serotonin, and norepinephrine cell bodies, please see Batten et al., 2025, Figure S1.

### In vivo data acquisition

A carbon fiber electrode manufactured in accordance with Kishida et al., 2016 and Batten et al. 2024 was inserted into SCC using a guide cannula which was part of the DBS procedure. The electrode was modified for an Alpha Omega microelectrode recording setup, with an impedance of 20 kΩ (DBS RC+S = 0.25 - 2 kΩ; PC = 0.35 - 8 kΩ). During the task, a standard triangle voltage waveform was applied to the electrode at 10Hz. The waveform voltage profile was: ramp at 400 V/s from -0.6V to +1.4V, back down from +1.4V to -0.6V (10ms), and then hold for 90ms at -0.6 V. The sampling rate for recording the resulting current was 100kHz. Electrode integrity was verified through characteristic current trace patterns during measurements and post-implantation microscopic inspection; no electrodes broke during the study.

### Machine learning-enhanced predictive modeling of in vivo neurochemical levels

Estimates of in vivo neurochemical concentrations were made by averaging the estimates of five neural network models applied to the in vivo current traces. Each of the five neural network models was identical in architecture (a modification of InceptionTime ^95^; see Batten et al., 2024, for full details of the model and training) and differed only due to the stochastic elements of the training (initial weights, train/validation split, and batch selection). Each ensemble model was trained using current-analyte concentration pairs obtained by exposing multiple (59) carbon electrodes to known concentrations of analytes. The model was originally validated using 6 test electrodes not used in the training data ^4^. **Extended Data Fig. 4a** shows the results of performing a full 10-fold cross validation of the model using 65 (59 + 6) electrodes. Each panel in **Extended Data Fig. 4b** corresponds to a specific analyte (DA, 5-HT, NA, pH). For each probe in the test fold we applied the model trained on the remaining 9 folds to produce points (x, y), with x being a true analyte value and y the associated estimated value; for a specific panel we took (x, y) pairs for the analyte under consideration and for the other analytes we took (x, e), where e is the error = estimate – true. For each of the analytes DA, 5-HT, and NA we then binned all of the x values into 28 equally spaced bins (and for pH, NN bins), and averaged either the estimated analyte values or error values for each probe. Then, for each bin we averaged the x values and either the average predictions or errors across probes and plotted these pairs. The error bars used probe as the degree of freedom. In training, the analyte concentration labels were transformed by z-scoring and then translating by 10. The model outputs were transformed back using the inverse transformation. For Supplementary Figure 1, the raw model outputs were used.

### Neurotransmitter signal processing and analysis

Sample-level neurotransmitter concentration estimates (DA, 5-HT, NA) were obtained from the machine learning model at 100 ms intervals. For the reversal learning task, each trial contained 101 samples; for the ultimatum game, each trial contained 61 samples.

For each participant, task, and neurotransmitter, all samples from both pre-stimulation and post-stimulation sessions were concatenated into a single time series. Each participant’s time series was then z-scored to standardize baseline neurotransmitter levels across participants while maintaining the relative magnitude of fluctuations within each participant’s data (**Extended Data Fig. 1**). Raw neurotransmitter data before z-scoring showing inter-trial and inter-participant variability are available in Supplementary Figure 1, and summary of participants’ neurochemical changes are presented in Supplementary Table 1.

Trial-level neurotransmitter estimates were computed by summing concentration values within a 500 ms window (5 consecutive samples) following task-relevant events. For the reversal learning task, the reference event was outcome reveal; for the ultimatum game, the reference event was offer presentation. Additional exploratory analyses of task-related features in relation to stimulation modulated neurotransmitter changes are provided in Supplementary Results and Tables 6–20.

Stimulation-induced changes in neurotransmitter levels were quantified by calculating participant-level difference scores: the mean trial-level estimate across all post-stimulation trials minus the mean trial-level estimate across all pre-stimulation trials for each neurotransmitter and task combination.

### Trial exclusions

When analyzing neurotransmitter estimates, we excluded trials where the response times were greater than 10 s to minimize the impact of distractions in the operating room (reversal learning: 0.8% of trials; ultimatum game: 0.8% of trials).

---

**Extended Data Fig. 1:**
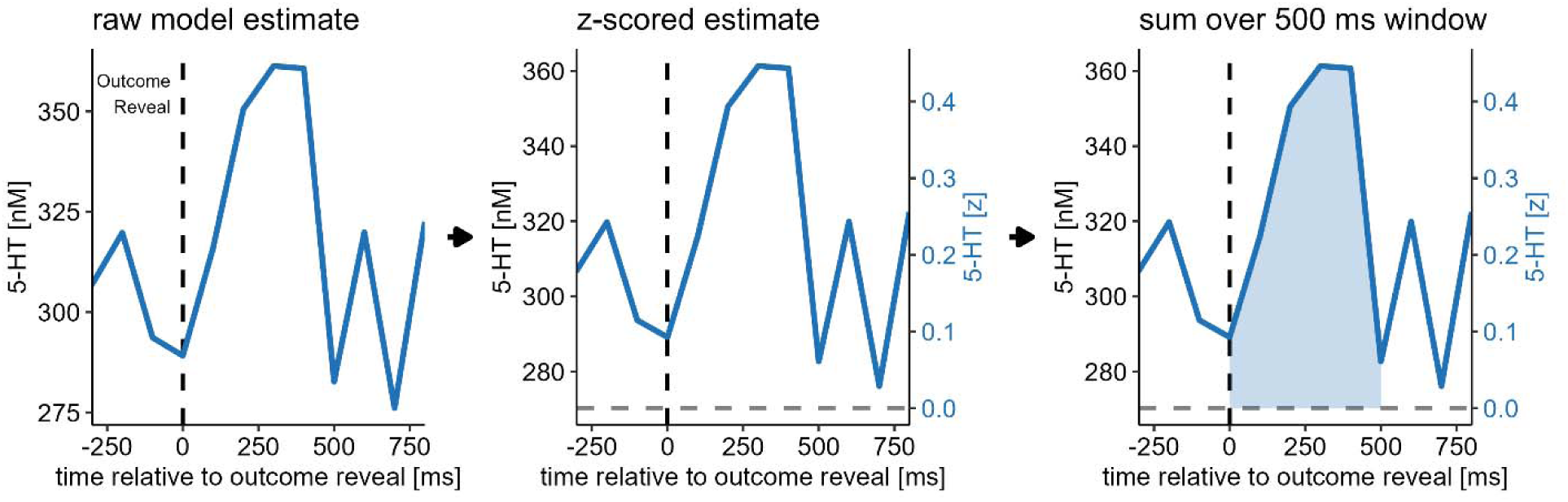
Analysis pipeline for in vivo neurotransmitter measurements. Representative example showing the computational steps used to transform raw machine learning model outputs into trial-level neurotransmitter estimates. Raw serotonin (5-HT) concentration estimates from the ensemble neural network model for a single trial during the reversal learning task (participant 5, post-stimulation session, reversal learning task, trial 28). Dashed vertical line indicates outcome reveal (time = 0). The same data after z-score transformation, which standardizes baseline neurotransmitter levels across participants while preserving within-participant temporal dynamics. Trial-level estimate computed by summing z-scored values within a 500 ms window (5 samples) following outcome reveal (shaded area). This windowing approach captures the phasic neurotransmitter response to task events while reducing noise from individual time points. The dual y-axes show both the original nanomolar scale (left) and z-score units (right) to illustrate the transformation. Arrows indicate the sequential processing steps applied to all participants, tasks, trials, and neurotransmitters (e.g., DA, 5-HT).

**Extended Data Fig. 2:**
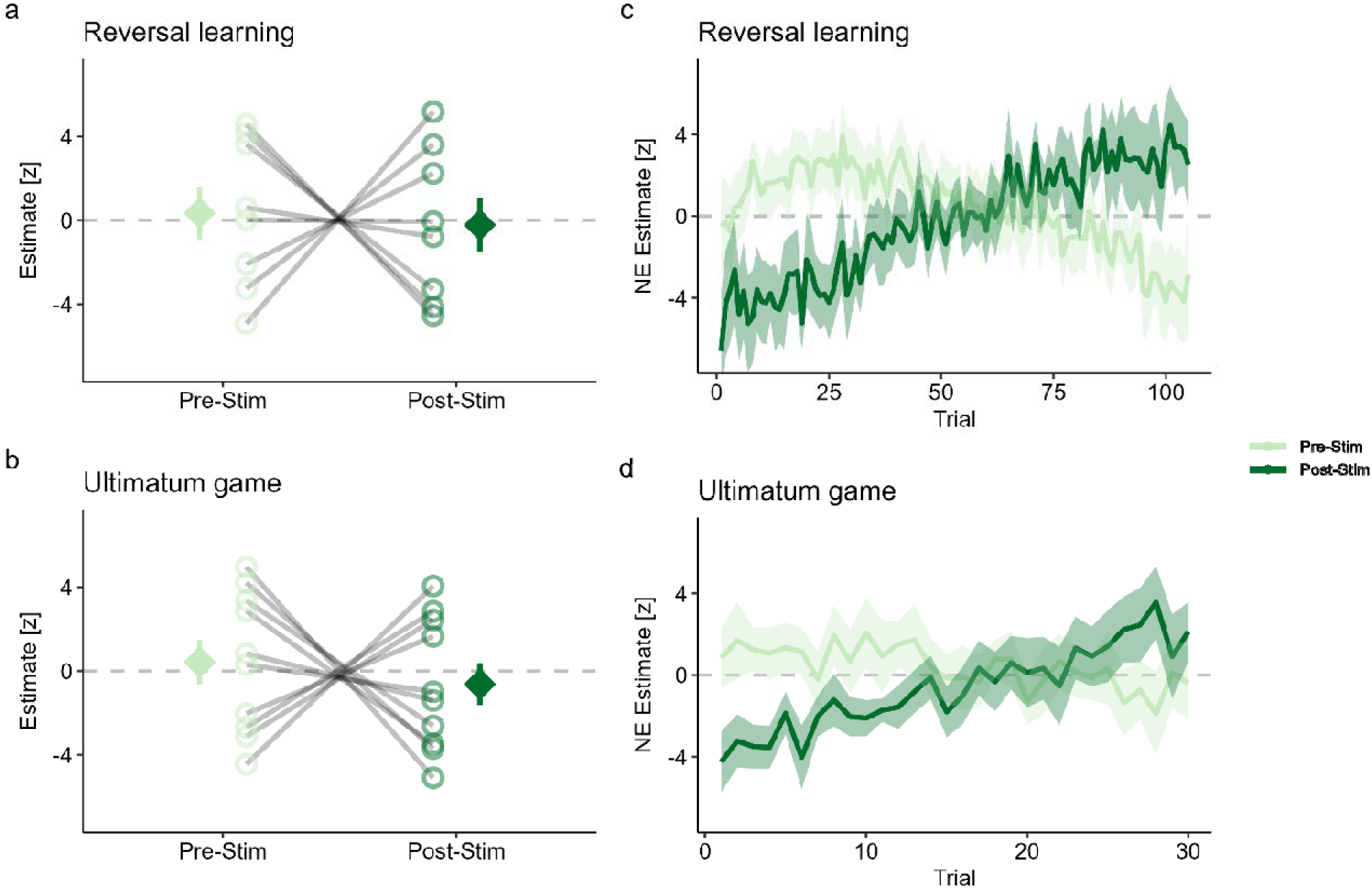
Norepinephrine levels remain unchanged following acute SCC stimulation. **a-b**, Stimulation-induced changes in norepinephrine (NE) estimates during the reversal learning (RL) task. Unlike DA and 5-HT, post-stimulation changes in NE following stimulation (repeated-samples *t* = 0.22, P = 0.832). **c-d**, Similarly, post-stimulation NE levels during the ultimatum game (UG) were unchanged (*t* = 0.51, P = 0.621). Error bars and shaded regions represent standard error of the mean across participants.

**Extended Data Fig. 3:**
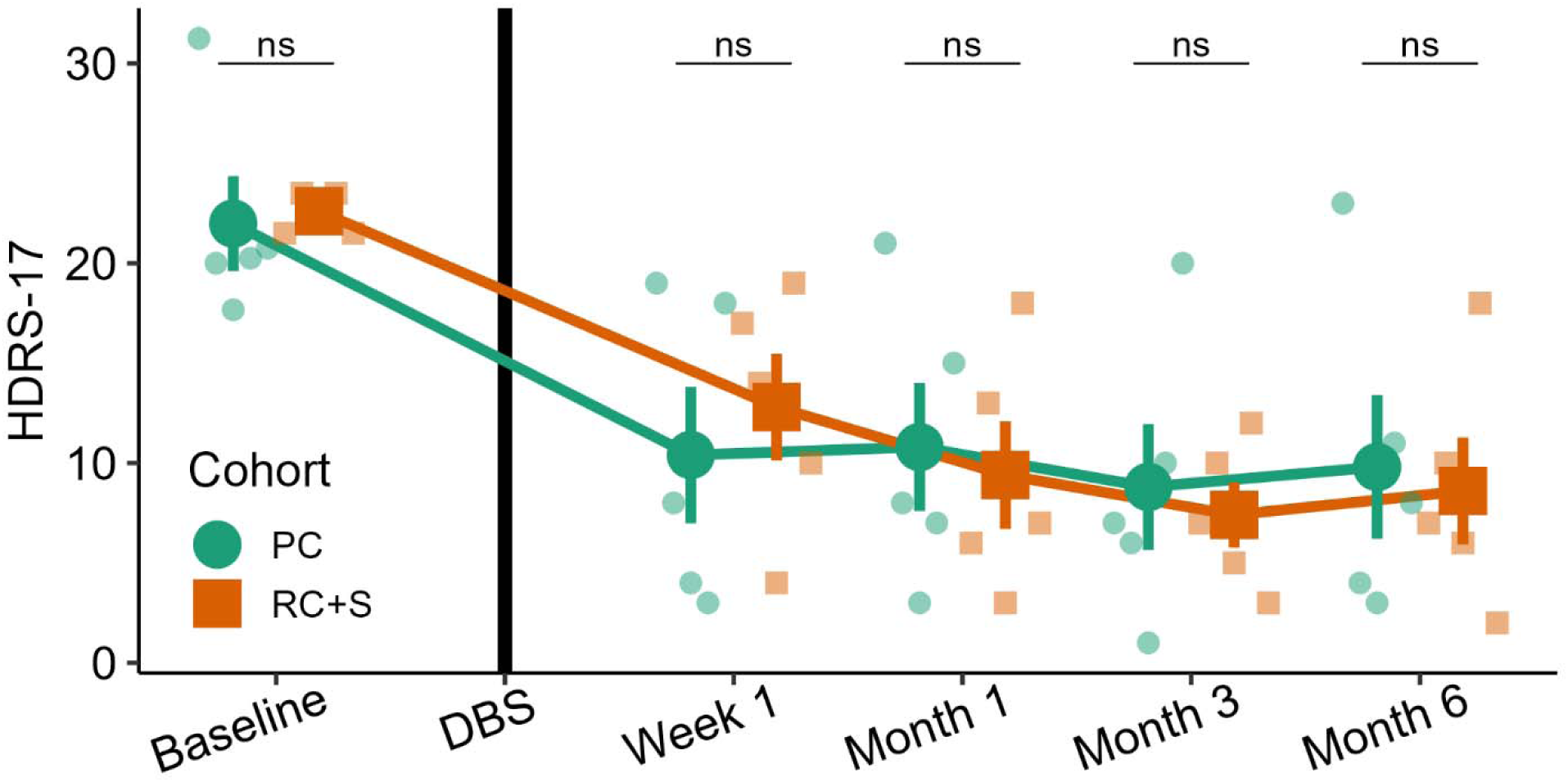
HDRS timeline of clinical cohorts. Trajectories of HDRS-17 scores over six months for 10 participants, split by clinical cohort. Cohorts received either the Medtronic PerceptPC or Medtronic Summit RC+S device. While the PC cohort had their devices turned on one day after implantation surgery, the RC+S cohort had their devices turned on one month after surgery. Smaller shapes indicate individuals while the larger shapes and lines indicate the mean, with error bars indicating standard error of the mean across participants. Statistical tests from two-tailed t-tests.

**Extended Data Fig. 4:**
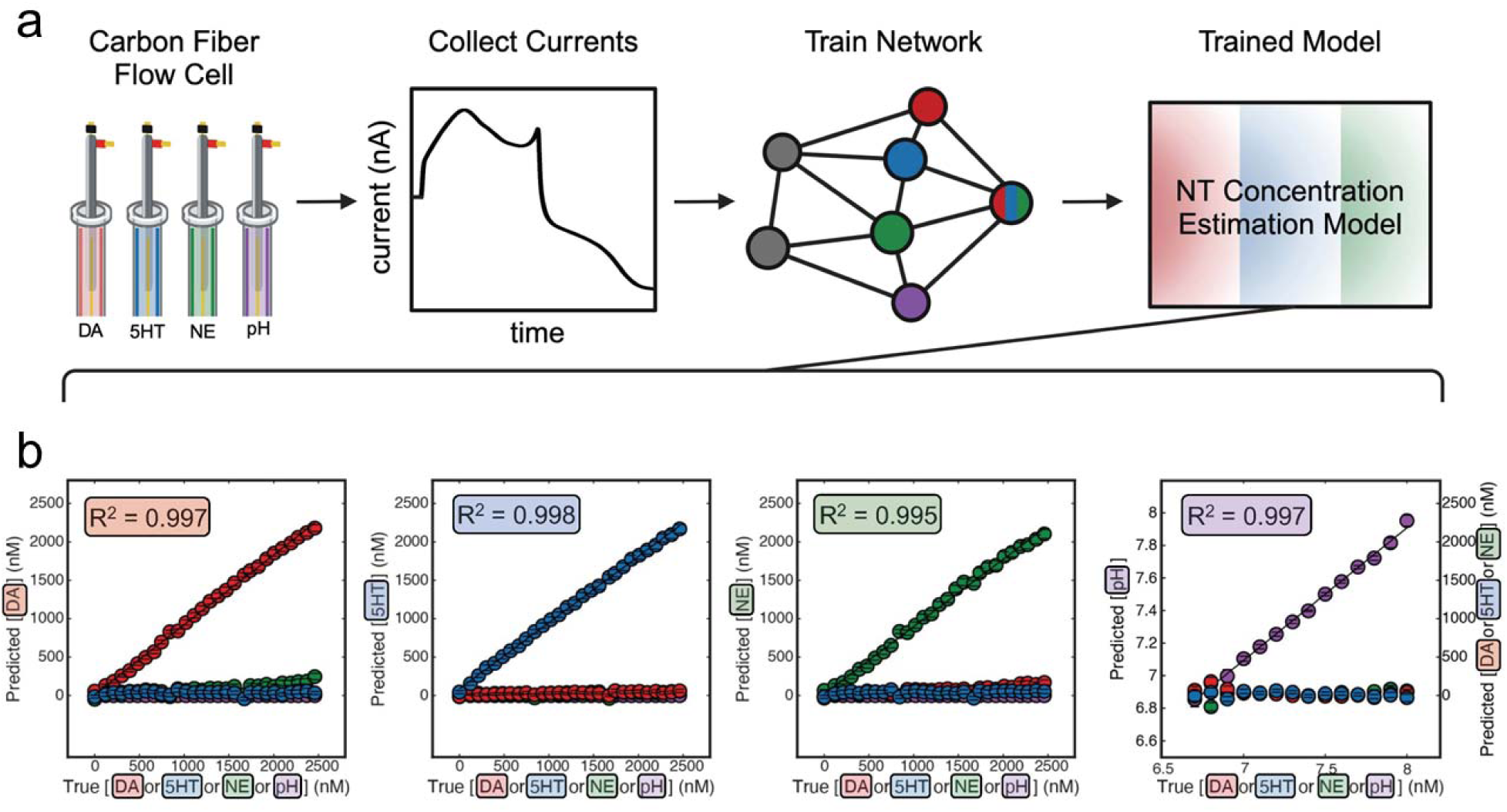
In vitro training and validation of neurochemical concentration estimation model. **a,** Training pipeline. 65 carbon fiber electrodes were exposed to known concentrations of dopamine (DA), serotonin (5-HT), norepinephrine (NE), and controlled pH values. The recorded voltammetric currents and their corresponding concentration labels were used to train a neural network model that generalizes to probes not included in the training set. **b,** 10-fold cross-validation plots. For each analyte (DA, 5-HT, NE, and pH), the true concentration values were binned and predicted values were averaged within each probe. These within-probe averages were then averaged across all probes within each bin. The error bars represent variability across probes within each bin, using probes as the degree of freedom. The solid line shows the linear fit between the binned average true concentrations and the corresponding average predicted concentrations, with the R^2^ value indicating the goodness-of-fit of this line.

**Extended Data Table 1.**
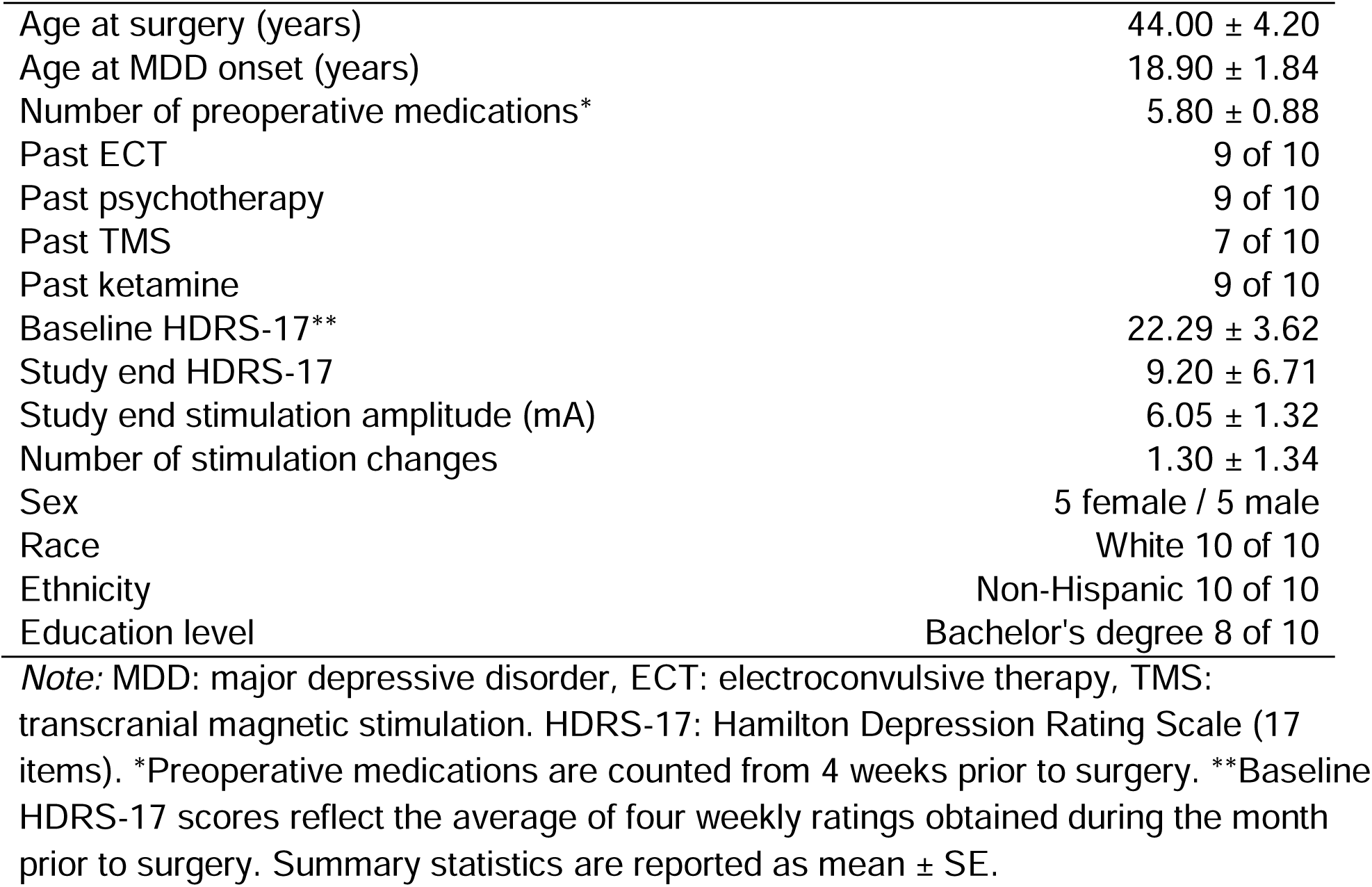
Overview of study participants’ demographics.

**Extended Data Table 2.** Summary of study participants’ stimulation parameters.

| Participant | HDRS-17 |  | Stimulation Amplitude (mA) |  |
| --- | --- | --- | --- | --- |
|  | Baseline | Study End | Study Start | Study End |
| 1 ** | 21.50 | 7.00 | 4.50 | 7.00 |
| 2 * | 23.50 | 10.00 | 4.50 | 6.00 |
| 3 ** | 23.00 | 6.00 | 4.50 | 4.50 |
| 4 # | 23.50 | 18.00 | 4.50 | 6.00 |
| 5 ** | 21.50 | 2.00 | 4.50 | 4.50 |
| 6 # | 31.25 | 23.00 | 4.50 | 8.00 |
| 7 ** | 20.00 | 4.00 | 4.50 | 6.00 |
| 8 ** | 17.67 | 3.00 | 4.50 | 6.00 |
| 9 # | 20.25 | 11.00 | 4.50 | 4.50 |
| 10 * | 20.75 | 8.00 | 4.50 | 8.00 |

*Note:* HDRS-17: Hamilton Depression Rating Scale (17 items). Participants are listed in chronological order of implantation with either the Medtronic Summit RC+S (participant 1-5) or Percept PC devices (participant 6-10). SCC DBS treatment outcomes are indicated as follows: \* responders (n = 7), \*\* remitters (n = 5), and # non-responders (n = 3). A 'response' was defined as a decrease in HDRS-17 scores greater than 50% of the presurgical baseline. Remission was defined as HDRS-17 < 8. Baseline HDRS-17 scores reflect the average of four weekly ratings obtained during the month prior to surgery. High-frequency stimulation was delivered at 130 Hz for participants implanted with RC+S device and 125 Hz for the Percept PC device.

## Acknowledgments

The authors are grateful to the patients and their families for their collaboration that enabled this study. The authors thank Gutemberg (Bobby) Forestal, Aida Razavilar, and Maryam Khalid for assistance with longitudinal data collection. Jessica Alexander and Laureen Pagan are acknowledged for their support with clinical administration and DBS patient coordination. The Summit RC+S and Percept PC devices were provided by Medtronic. The authors also thank the surgical team and the Alpha Omega technical team at Mount Sinai West for their support.

This work was supported by the National Institute of Health grants R01MH124115 (XG, PRM), UH3NS103550 (HSM), the Hope for Depression Foundation (HSM), and the Red Gates Foundation (PRM). Work on “Computational Roles of Dopamine and Serotonin Transients in Anhedonia in Humans with Treatment Resistant Depression“ was supported by Wellcome Leap as part of the Multi-Channel Psych Program (XG,HSM).

This work was supported in part through the computational and data resources and staff expertise provided by Scientific Computing and Data at the Icahn School of Medicine at Mount Sinai and supported by the Clinical and Translational Science Awards (CTSA) grant UL1TR004419 from the National Center for Advancing Translational Sciences.

The content is solely the responsibility of the authors and does not necessarily represent the official views of the funders.

## Code availability

All analysis code and scripts for visualization is publicly available on GitHub at https://github.com/blairshevlin/shevlin-fu_volt-trd.

## Data availability

De-identified data will be made publicly available upon publication on GitHub at https://github.com/blairshevlin/shevlin-fu_volt-trd.

## Contributions

XG, HSM, and PRM led the conceptualization, with support from BRKS, MF, RM, DB, VGF, and KTK. Methodology was led by XG, PRM, and WMH, with support from BRKS, SRB, SH, CJR, AK, KRK, KSC, HNS, JPW, TL, MF, RM, DB, KTK, LSB, IS, and HSM.

Software was developed by PRM, QXF, LSB, and TT, with support from BRKS, MH, SH, KRK, JPW, TL, RM, WMH, and XG. Validation was led by PRM, with support from QXF, SRB, MH, AK, AEH, WMH, KTK, LSB, and HSM.

QXF, AND, MH, and XG led the investigation, supported by SRB, BHK, SH, AK, IT, MF, KTK, IS, and HSM. Clinical expertise was provided by HSM and MF, with support from BHK, KSC, SO, and IS. Resources were coordinated by QXF, HSM, and PRM, with support from SRB, MH, SH, AK, KRK, SO, TN, IT, JPW, RM, WMH, VGF, and XG. Data curation was led by QXF and PRM, with support from BRKS, SRB, MH, SH, AK, KSC, TN, JPW, MF, TT, RM, LSB, HSM, and XG.

Formal analysis was led by BRKS and PRM, supported by QXF, SB, CJR, KRK, DB, LSB, HSM, and XG. BRKS and QXF led the original draft writing, with support from XG and HSM. Review and editing were led by BRKS, QXF, and XG, with support from BHK, MH, CJR, AK, SO, TL, MF, DB, WMH, KTK, LSB, VGF, IS, PRM, and HSM.

Visualization was led by BRKS, with support from QXF, AK, HNS, AEH, WMH, PRM, and XG.

Supervision was provided by XG and PRM, with support from HSM, BRKS, MF, and WMH. Project administration was led by QXF, MH, HSM, and XG, with support from SH, TN, IT, and PRM.

