## Supplementary Information for "Dopamine and serotonin transients predict depressive symptom relief following deep brain stimulation of human subcallosal cingulate cortex"

### Supplementary Results

#### Temporal dynamics analysis

To assess whether neurotransmitter changes reflected learning/repetition effects versus stimulation-specific effects, we examined how neurotransmitter estimates changed across trials within each task session. We fit linear mixed-effects models with a three-way interaction between stimulation condition (pre/post), task (reversal learning/ultimatum game), and standardized trial number, with random intercepts and slopes for stimulation and task by participant.

Simple slopes were extracted using the *emtrends* function from the *emmeans* package in R to quantify the rate of change (slope) in neurotransmitter levels per trial for each neurotransmitter system, task, and stimulation condition.

For visualization (Supplementary Figure 2), we compared neurotransmitter estimates between early trials (first quartile) and late trials (fourth quartile) within each task session. Panel a displays these temporal slopes with 95% confidence intervals. Panels b-c show individual participant trajectories (thin transparent lines) and group means ± SEM (thick lines) comparing early versus late trial periods within each task and stimulation condition.

#### Sensitivity analysis of neurotransmitter quantification

To assess the robustness of our approach for quantifying neurotransmitter levels, we tested alternative metrics beyond the sum-based estimate reported in the main text. We evaluated four metrics at two temporal windows (500 ms and 1000 ms following task-relevant events): (1) overall estimate: sum of concentration values (main text metric); (2) mean estimate: average concentration within the window; (3) peak estimate: maximum concentration within the window; and (4) relative estimate: sum corrected by subtracting the baseline value at event onset.

For each combination of neurotransmitter, task, metric, and window, we conducted two-way repeated-measures ANOVAs testing for session effects (pre- vs. post-stimulation), task feature effects (RL: block type; UG: offer amount), and their interactions.

Results confirmed our main findings across most metrics (Supplementary Table 23). In the reversal learning task, the overall, mean, and peak estimates all detected increased DA following stimulation (all Ps < 0.001) regardless of window duration, while no metric detected 5-HT changes (all Ps > 0.05). In the ultimatum game, these same three metrics confirmed post-stimulation increases in both DA and 5-HT across both windows (all Ps < 0.05).

The relative estimate failed to detect stimulation-induced changes in either task. This was expected, as this metric quantifies change relative to local baseline rather than absolute concentration levels, consistent with our interpretation that stimulation altered overall neurotransmitter availability rather than task-specific functional dynamics in the SCC.

#### Alternative normalization of neurotransmitter estimates

To address potential concerns that our primary z-scoring approach might introduce systematic biases by normalizing across tasks with different temporal structures, we implemented an alternative, more conservative normalization procedure. This approach jointly z-scored neurotransmitter estimates from both tasks using only temporally matched event windows, ensuring that any observed differences between tasks or stimulation conditions could not arise from task-specific normalization artifacts.

Rather than z-scoring across all samples from an entire experimental session (as in the primary analysis), we restricted normalization to a narrow temporal window around task-relevant events. We extracted neurotransmitter concentration estimates within ±1 second of task-relevant events (±10 samples at 10 Hz sampling rate). For the ultimatum game, this corresponded to samples 20-41 relative to offer presentation; for reversal learning, this corresponded to samples 40-61 relative to outcome reveal. We aligned comparable events across tasks. For reversal learning, this was the outcome reveal (when participants received reward/punishment feedback); for the ultimatum game, this was the offer presentation (when participants received the monetary offer). These events were designated as "Shared" events for cross-task normalization, as both represent the presentation of outcome-relevant information that drives subsequent decision-making.

For each participant and neurotransmitter, we concatenated the restricted time-window samples from both tasks and both sessions (pre- and post-stimulation) into a single time series. For reversal learning this was: 22 samples/trial × 105 trials × 2 sessions = 4,620 samples. For the ultimatum game this was: 22 samples/trial × 30 trials × 2 sessions = 1,320 samples.

This concatenated time series was then z-scored. This procedure ensures that neurotransmitter estimates are on the same scale across both tasks; any differences between tasks reflect genuine differences in neurotransmitter dynamics, not normalization artifacts ; and within-participant variability is preserved while controlling for between-participant baseline differences.

Following z-scoring, we computed trial-level neurotransmitter estimates by averaging z-scored concentration values within a 500 ms window (5 consecutive samples) following task-relevant events, consistent with our primary analysis approach.

The alternative normalization procedure replicated our primary findings (Supplementary Figure 5). In the reversal learning task, there was a significant, post-stimulation increase in dopamine (F_1,42_ = 20.47, P < 0.001), but not serotonin (F_1,42_ = 0.82, P = 0.372). In the ultimatum game, there was a significant, post-stimulation increase in serotonin (F_1,54_ = 1.57, P = 0.216), but not dopamine (F_1,54_ = 25.25, P < 0.001).

We also tested for the task × session interactions. Both dopamine (F_1,32_ = 6.66, P = 0.015) and serotonin (F_1,32_ = 4.89, P = 0.034) showed significant main effects of stimulation. Critically, the task × stimulation interaction was non-significant for both neurotransmitters (dopamine: F_1,32_ = 3.07, P = 0.089; serotonin: F_1,32_ = 1.27, P = 0.269), indicating that while the magnitude of effects may differ between tasks, the direction of stimulation effects was consistent.

#### Analysis of trial-level neural data

Linear mixed-effects regressions assessed trial-level neural data within each task with random effects at the participant level. Models were computed using the *lmer* function from the *lme4* and *lmertest* packages (Bates et al., 2015; Kuznetsova et al., 2017) in R v4.4.3 (R Team, 2025). Significant main effects and interactions were probed by computing estimated marginal means using the *emmeans* function from the *emmeans* R package (Leanth & Piaskowski, 2025).

Across tasks, session was binary coded (-1 = pre-stimulation session, 1 = post-stimulation session), response times were log-transformed, and trial numbers were z-score normalized within session and participant.

In the reversal learning task, the following conventions were used: outcome (0 = loss, 1 = win), error trial (0 = suboptimal chosen, 1 = optimal chosen), post-reversal (1 = within five trials of a reversal, 0 = otherwise), slow trial (0 = response time less than or equal to the participant -level median, 1 = response time greater than the participant-level median). Block type was dummy coded with the Mixed level as the reference category (Negative: 1, 0; Positive: 0, 1; Mixed: 0, 0). Both current reward and previous reward were dummy coded with an outcome of $0 as the reference category (Lose $10: 1, 0; Win $10: 0, 1: Lose/Win $0: 0, 0). Prediction error was dummy coded with No PE (current reward equal to previous reward) as the reference category (Negative PE: 1, 0: Positive PE: 0, 1: No PE: 0, 0).

In the ultimatum game task, the following conventions were used: offer amount was z-score normalized, accept trial (0 = offer rejected; 1 = offer accepted), slow trial (0 = response time less than or equal to the participant -level median, 1 = response time greater than the participant-level median). Both current offer and previous offer were dummy coded with offers $4-6 as the reference category (Low/$1-3: 1, 0; Middle/$7-9: 0,1: High/$4-6: 0, 0). Offer change was dummy coded with no change (current offer equal to previous offer) as the reference category (offer worsened: 1, 0: offer improved: 0, 1: no change: 0, 0).

In Wilkinson notation, the linear neural models (N) for the reveral learning task (RL) and ultimatum game (UG) were specified as:

- N-RL-M1, estimate_DA/5-HT/NE_ ~ 1 + session × (outcome + block type + trial number) + (1 + stimulation | participant)
- N-RL-M2, estimate_DA/5-HT/NE_ ~ 1 + session × (error trial + post reversal + slow trial + trial number) + (1 + stimulation | participant)
- N-RL-M3, estimate_DA/5-HT/NE_ ~ 1 + session × (current reward × previous reward + trial number) + (1 + stimulation | participant)
- N-RL-M4, estimate_DA/5-HT/NE_ ~ 1 + session × (prediction error + trial number) + (1 + stimulation | participant)
- N-UG-M1, estimate_DA/5-HT/NE_ ~ 1 + session × (offer amount + trial number) + (1 + stimulation | participant)
- N-UG-M2, estimate_DA/5-HT/NE_ ~ 1 + session × (accept trial + slow trial + trial number) + (1 + stimulation | participant)
- N-UG-M3, estimate_DA/5-HT/NE_ ~ 1 + session × (current offer × previous offer + trial number) + (1 + stimulation | participant)
- N-UG-M4, estimate_DA/5-HT/NE_ ~ 1 + session × (offer change + trial number) + (1 + stimulation | participant).

For the reversal learning task, we found a significant interaction between stimulation session and block type (Supplementary Table 6; Mixed – Positive: *β* = -0.55, *t* = -2.31, P = 0.021) on 5-HT estimates and a significant effect of block type (Mixed – Positive: *β* = 0.89, *t* = 2.48, P = 0.013) on NE estimates. However, follow-up pairwise comparisons of the estimated marginal means were non-significant except for the contrast between Negative and Positive blocks on NE estimates in the pre-stimulation session (estimate = -1.21, *t* = 2.36, P = 0.048).

We found a significant interaction between stimulation session and error trials on 5-HT estimates (Supplementary Table 7; *β* = 0.26, *t* = 2.05, P = 0.041). However, pairwise comparisons of estimated marginal means were non-significant (Ps > 0.05).

We found significant effects of the interaction between stimulation session and current losses (Supplementary Table 8; *β* = -0.61, *t* = -2.16, P = 0.031) and the three-way interaction between stimulation session, current losses, and previous losses (*β* = 0.87, *t* = 1.98, P = 0.048) on DA estimates. Follow-up pairwise comparisons of the estimated marginal means show two significant contrasts, both following Win/Loss of $0 during the pre-stimulation session (Win/Lose $0 – Lose $10 estimate = -1.01, *t* = -2.48, P = 0.035; Lose $10 – Win $10 estimate: 1.53, *t* = 3.01, P = 0.008).

We found a significant effect of prediction error type (Supplementary Table 9; No PE – Positive PE: *β* = -0.46, *t* = -2.15, P = 0.032) on NE estimates. Follow-up pairwise comparisons of the estimated marginal means were non-significant (Ps > 0.05).

For the ultimatum game, we found significant effects of offer amount (Supplementary Table 10; *β* = 0.95, *t* = 2.04, P = 0.042) and its interaction with stimulation session (*β* = -1.39, *t* = -2.12, P = 0.035) on DA estimates. Pairwise comparisons of the estimated marginal means revealed high offers were associated with increased DA estimates compared to low offers in the pre-stimulation session (Low – High estimate = -0.95, *t* = -2.03, P = 0.044), but not the post-stimulation session (Low – High estimate = 0.44, *t* = 0.95, P = 0.344).

Acceptance rates and its interactions with stimulation session also did not have a significant relationship with neurotransmitters estimates (Supplementary Table 11, Ps > 0.05). However, we did observe as significant interaction between slow trials (*β* = -0.82, *t* = -2.86, P = 0.004) and its interaction with stimulation session on 5-HT estimates (*β* = 1.21, *t* = 3.03, P = 0.003). However, the pairwise comparisons of the estimated marginal means did not reveal significant differences in fast and slow trials’ 5-HT estimates during either stimulation session (Ps > 0.05).

NE estimates were found to vary based on previous offer amount (Supplementary Table 12, Middle - High: *β* = -2.06, *t* = -2.01, P = 0.045), the interaction between session and previous offer amount (*β* = 3.10, *t* = 2.21, P = 0.028), and the interaction between session, current offer amount, and previous offer amount (*β* = -4.57, *t* = -2.29, P = 0.023). Using pairwise comparisons, the only significant contrast was between Low and High offers when the previous offer was Low in the pre-stimulation session (estimate = 2.02, *t* = 2.06, P = 0.040).

No significant relationships between offer changes and neurotransmitter estimates were observed (Supplementary Table 13).

### Supplementary Figures

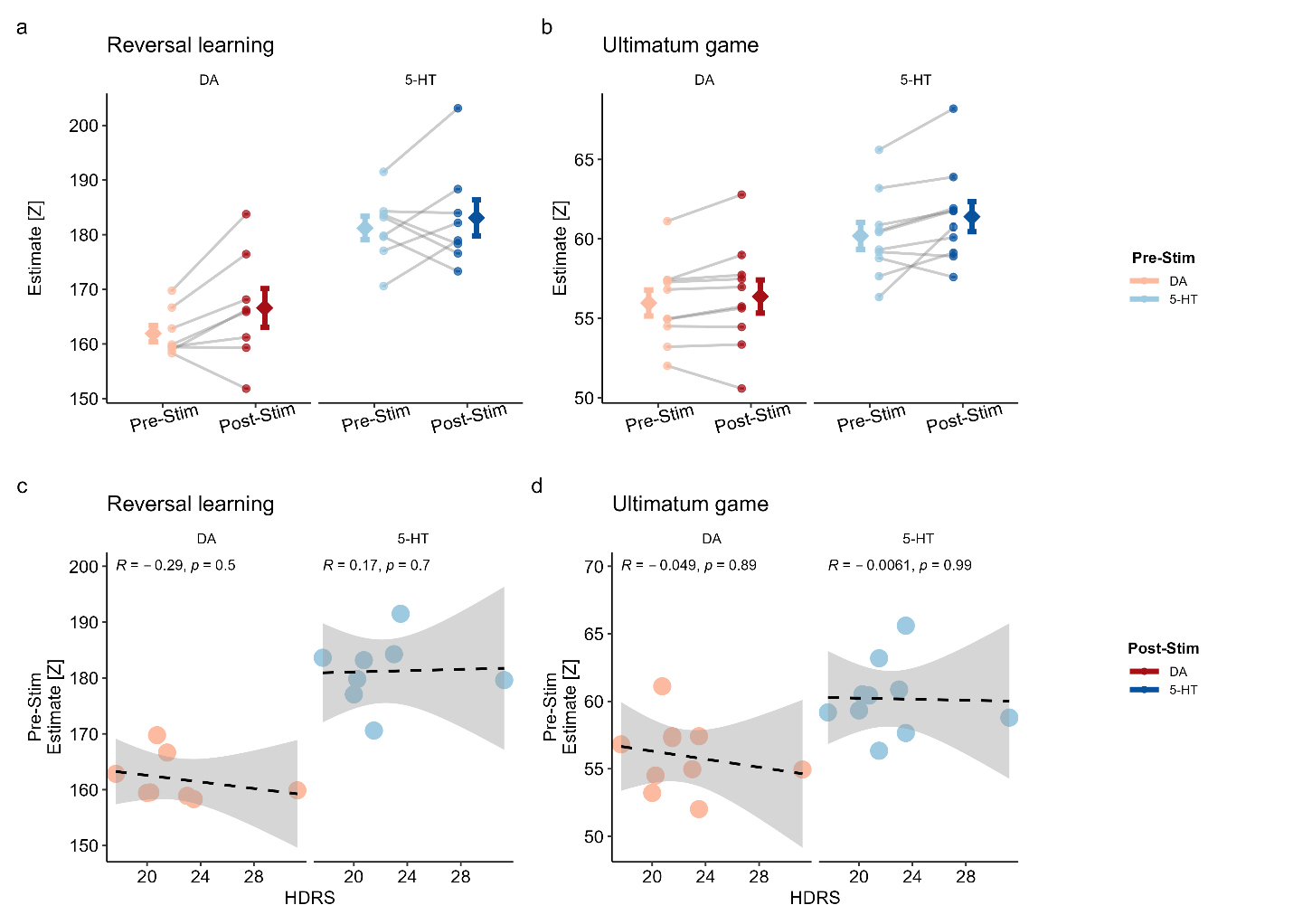

#### **Supplementary Figure 1. Raw neurotransmitter data before across-session z-scoring showing inter-trial and inter-participant variability.**

**a-b,** Individual participant mean estimates (circles) with error bars indicating inter-trial variability (SEM) for dopamine (DA, red) and serotonin (5-HT, blue) during **(a)** reversal learning and **(b)** ultimatum game tasks. Gray lines connect pre- and post-stimulation measurements for each participant. Group means are shown as diamonds with error bars representing inter-patient variability (SEM). Light colors indicate pre-stimulation; dark colors indicate post-stimulation. **c-d,** Correlations between individual participant baseline neurotransmitter estimates (pre-stimulation) and Hamilton Depression Rating Scale (HDRS) scores for **(c)** reversal learning **(d)** and ultimatum game tasks. Dashed lines show linear fits with 95% confidence intervals (gray shading). All correlations were non-significant (P > 0.05). Note that model estimated neurotransmitter values are in z-scored standardized units.

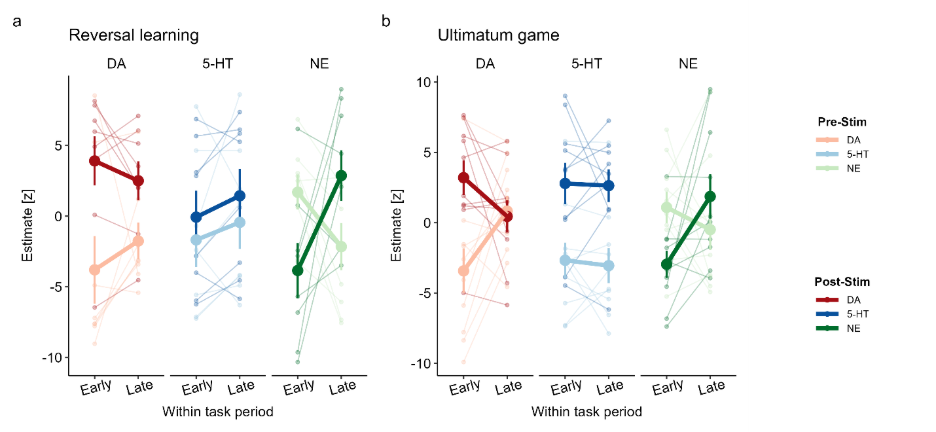

#### **Supplementary Figure 2. Task-specific temporal dynamics of neurotransmitter levels argue against learning effects.**

**a-b**, Individual participant trajectories comparing early (first quartile) versus late (fourth quartile) trials within the reversal learning task (**a**) and ultimatum game (**b**). If the observed between-session neurotransmitter changes reflected non-specific time-on-task effects (e.g., learning, repetition, arousal, etc.), within-session patterns should consistently parallel between-session effects. Linear mixed-effects models examined the three-way interaction between stimulation condition (pre/post), task, and standardized trial number, with random intercepts and slopes by participant. **a**, During reversal learning, within-session dopamine (DA) *increased* in the pre-stimulation session (t = -6.32, P < 0.0001) but *decreased* in the post-stimulation session (t = 4.33, P < 0.0001). This reversal of within-session DA dynamics is inconsistent with non-specific temporal effects and instead suggests stimulation altered the fundamental DA response pattern. Serotonin (5-HT) increased within both sessions (pre: t = -4.74, P < 0.0001; post: t = -6.21, P < 0.001), while norepinephrine (NE) decreased pre-stimulation (t = 9.60, P < 0.0001) and increased post-stimulation (t = -16.77, P < 0.001). **b**, During the ultimatum game, within-session DA again showed opposing patterns (pre-stim increase: t = -4.27, P < 0.0001; post-stim decrease: t = 5.08, P < 0.0001). Critically, 5-HT showed *no significant within-session changes* in either the pre-stimulation (t = 0.94, P = 0.348) or post-stimulation (t = 0.44, P = 0.661) sessions, despite showing robust between-session increases. This dissociation between flat within-session and elevated between-session 5-HT argues against non-specific temporal effects. NE decreased pre-stimulation (t = 2.88, P = 0.004) and increased post-stimulation (t = -8.67, P < 0.0001). The divergent patterns between tasks, combined with inconsistent within-session trajectories, indicate that the observed stimulation effects reflect genuine task-specific neuromodulation rather than non-specific time-on-task effects. Light colors: pre-stimulation; dark colors: post-stimulation. DA: dopamine (red/pink), 5-HT: serotonin (blue), NE: norepinephrine (green). Error bars represent 95% confidence intervals. Lines show individual participants (transparent) and group means (thick lines) ± SEM.

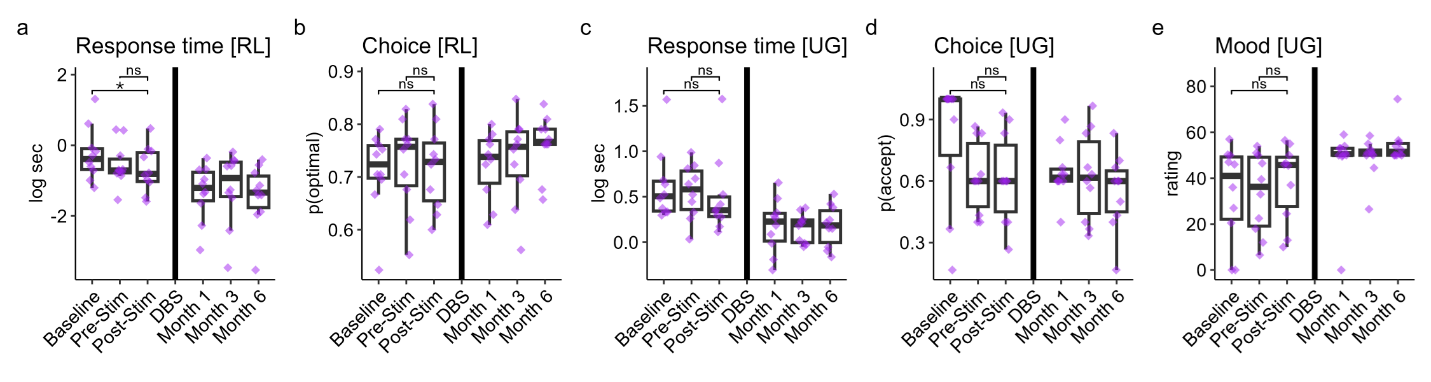

#### **Supplementary Figure 3. Intraoperative behavioral trends**.

**a,** Compared to baseline, participants’ response times during the reversal learning (RL) task were significantly reduced following acute simulation (paired-samples Wilcoxson test, P = 0.049). However, post-stimulation response times were not significantly different from pre-stimulation response times (P > 0.05). **b,** Optimal choice behavior during the RL task following acute stimulation did not significantly change compared to baseline or pre-stimulation sessions (Ps > 0.05). **c-e,** Behavioral patterns during the ultimatum game (UG) task following acute stimulation did not significantly change compared to baseline or pre-stimulation sessions (Ps > 0.05).

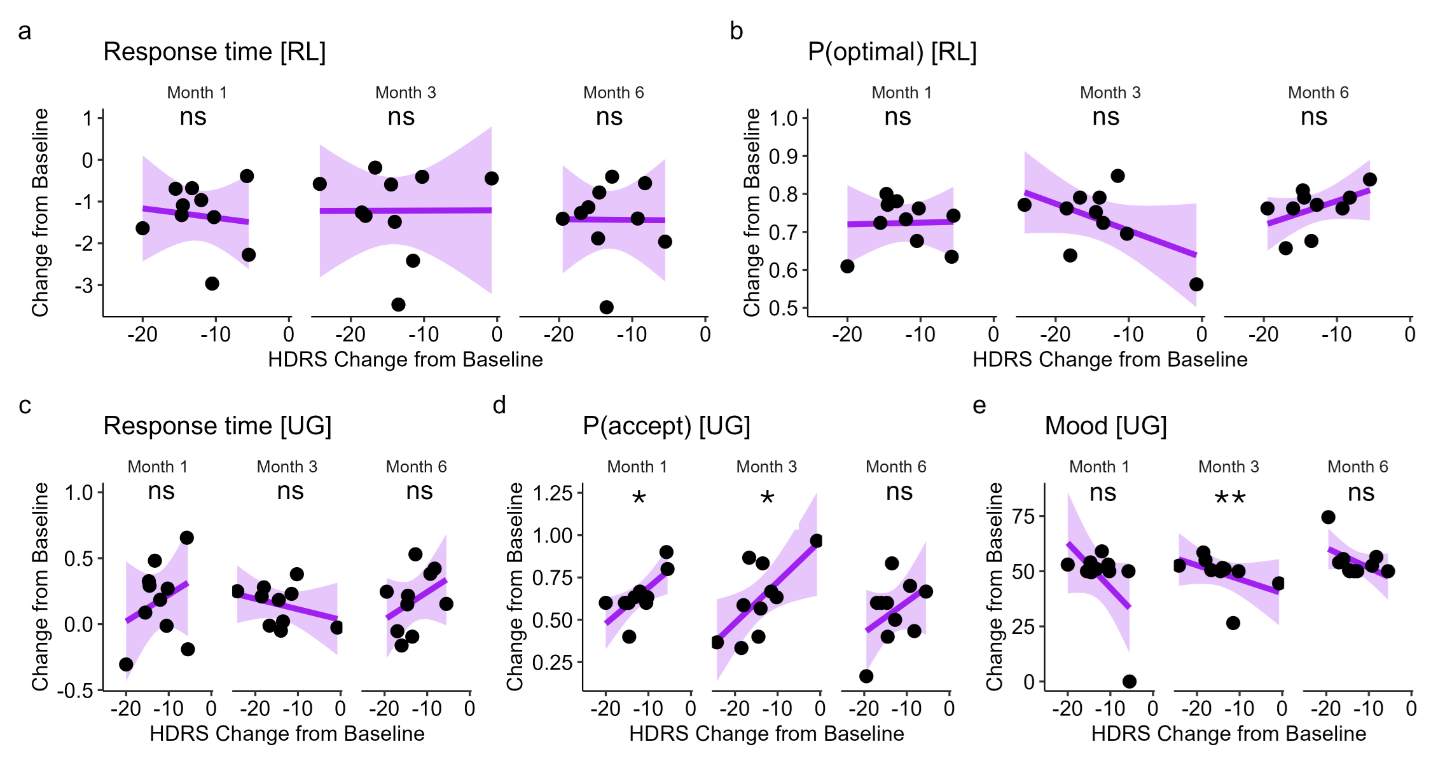

#### **Supplementary Figure 4. Associations behavioral longitudinal behavioral trends and symptom severity**.

**a-b,** Longitudinal changes in RL task performance were uncorrelated to changes in symptom severity, as measured by the Hamilton Depression Rating Scale (HDRS, Ps > 0.05). **c,** Longitudinal changes in response times during the UG task were uncorrelated with changes in HDRS scores (P > 0.05). **d,** After chronic DBS stimulation, decreases in offer acceptance rates during the UG task predicted greater decreases in HDRS scores in Month 1 (Spearman’s rho = 0.75, P = 0.013) and Month 3 (rho = 0.70, P = 0.031). **e,** Larger increases in mood ratings during the UG task at Month 3 predicted greater decreases in HDRS scores (rho = -0.81, P = 0.005).

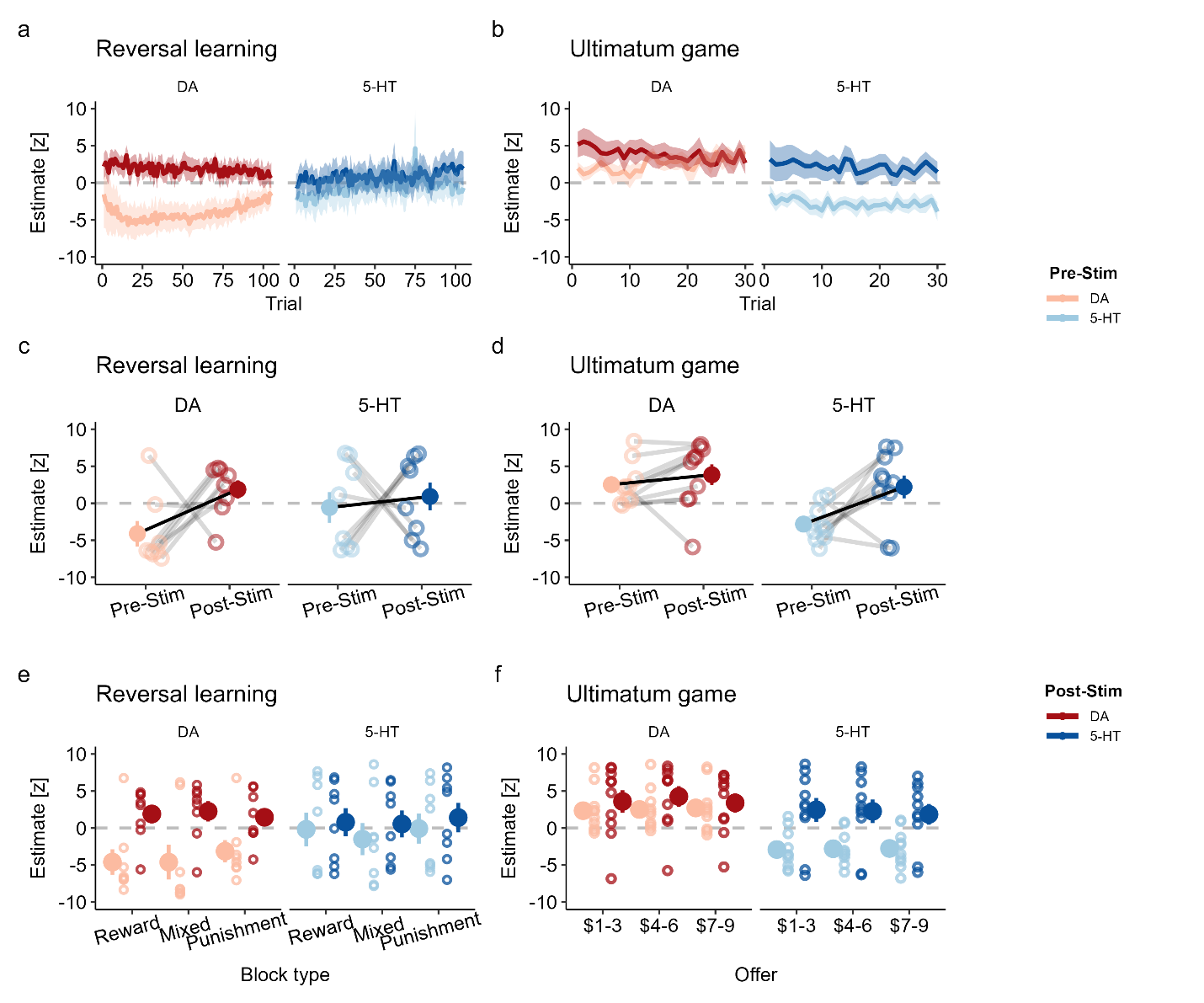

#### **Supplementary Figure 5. Alternative z-scoring normalization validates primary findings of stimulation-induced neurotransmitter modulation.**

**a-b,** Trial-by-trial mean neurotransmitter estimates across tasks. Solid lines represent group means; shaded regions represent ±1 SEM. **a,** Reversal learning demonstrates sustained dopamine elevation post-stimulation (dark red) compared to pre-stimulation (light red), with minimal change in serotonin. **b,** Ultimatum game demonstrates sustained serotonin elevation post-stimulation (dark blue) compared to pre-stimulation (light blue), with minimal change in dopamine. **c-d,** Participant-level mean estimates (z-scored) across pre-stimulation and post-stimulation sessions. Lines connect measurements within individual participants. Diamonds represent group means. **c,** Reversal learning shows significant increase in dopamine (DA) following stimulation (F₁,₄₂ = 20.47, P < 0.001) but no significant change in serotonin (5-HT) (F₁,₄₂ = 0.82, P = 0.372). **d,** Ultimatum game shows significant increase in serotonin following stimulation (F₁,₅₄ = 25.25, P < 0.001) but no significant change in dopamine (F₁,₅₄ = 1.57, P = 0.216). **e-f,** Task-specific feature modulation. Individual participant data points (circles) are shown with transparency; diamonds represent group means. **e,** Reversal learning neurotransmitter estimates by block type (Reward, Mixed, Punishment) show no significant interaction with stimulation (dopamine: F₂,₄₂ = 0.29, P = 0.753; serotonin: F₂,₄₂ = 0.04, P = 0.965). **f,** Ultimatum game neurotransmitter estimates by offer amount show no significant interaction with stimulation (dopamine: F₂,₅₄ = 0.10, P = 0.906; serotonin: F₂,₅₄ = 0.05, P = 0.953). These results replicate primary findings using a more conservative normalization approach that jointly z-scores both tasks, confirming that stimulation effects are robust to methodological choices in data preprocessing.

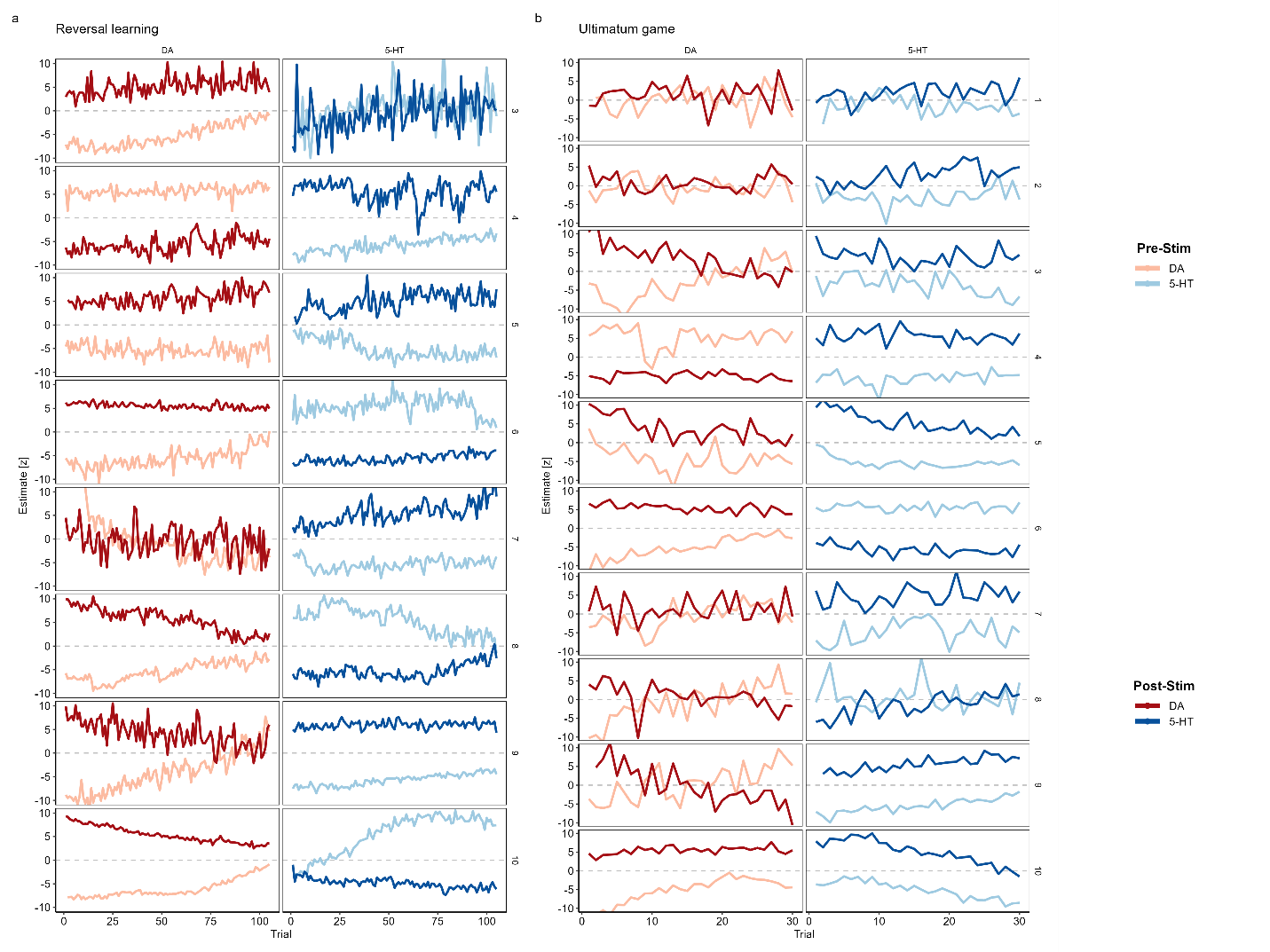

#### **Supplementary Figure 6. Individual participant neurotransmitter dynamics across behavioral tasks.**

Trial-by-trial dopamine (DA) and serotonin (5-HT) estimates for individual participants during reversal learning and ultimatum game tasks, before and after acute subcallosal cingulate (SCC) stimulation. Neurotransmitter estimates reflect z-scored concentration values summed within a 500 ms window following task-relevant events (outcome reveal for reversal learning; offer presentation for ultimatum game). **a,** Reversal learning task (105 trials) showing participant-specific DA and 5-HT dynamics. Each row represents an individual participant (n=8), with DA estimates in the left column and 5-HT estimates in the right column. Pre-stimulation estimates are shown in lighter colors (light red for DA, light blue for 5-HT), while post-stimulation estimates are shown in darker colors (dark red for DA, dark blue for 5-HT). **b,** Ultimatum game task (30 trials) showing the same participant-level organization and color scheme (n=10).

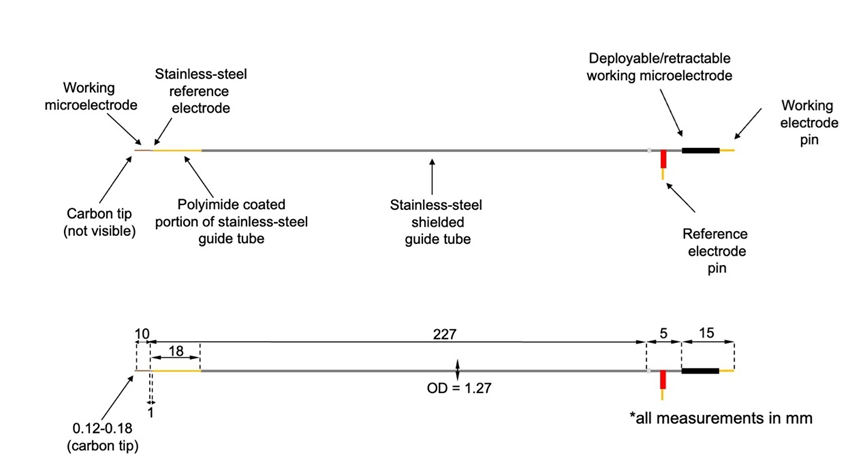

#### **Supplementary Figure 7. Dimensions of the human carbon fiber electrode used in the study.**

The electrode is approximately 250 mm in length. The carbon fiber electrode is made in polyimide coated silica and housed in an AlphaOmega guide tube (in order to fit other neurosurgical equipment and guide cannula). The carbon fiber measurement surface is 7µm in diameter and 120-180 µm in length. The carbon fiber is connected to a platinum-iridium wire within the silica via silver paint. The guide tube is 1.27 mm in diameter. The electrode is retracted and placed through a guide cannula until it reaches the correct dorsal-ventral placement in the brain. The carbon fiber electrode is then extended 10 mm into brain tissue to reach the measurement site and recordings are taken. The electrode is retracted back into the guide tube and removed from the guide cannula when the experimental recordings are over. For more details on the construction of the carbon fiber electrode, see Kishida et al., 2016.

### Supplementary Tables

#### **Supplementary Table 1. Summary of study participants’ neurochemical changes.**

Values shown are average z-standardized changes (post-stimulation minus pre-stimulation), with the standard error presented in parentheses. Reversal learning data for participants 1 and 2 are missing due to complications with timeline alignment.

| **Participant** | **Reversal learning** | | | **Ultimatum game** | | |
| --- | --- | --- | --- | --- | --- | --- |
|  | **ΔDA** | **Δ5-HT** | **ΔNE** | **ΔDA** | **Δ5-HT** | **ΔNE** |
| 1 | NA | NA | NA | 1.34  (0.55) | 3.39  (0.47) | 4.52  (0.39) |
| 2 | NA | NA | NA | 1.34  (0.42) | 5.83  (0.48) | -0.51  (0.47) |
| 3 | 10.44  (0.17) | -1.12  (0.33) | 10.12  (0.19) | 6.27  (1.04) | 7.57  (0.48) | 6.07  (1.05) |
| 4 | -11.31  (0.14) | 11.15  (0.21) | -9.30  (0.40) | -10.28  (0.39) | 11.24  (0.36) | -9.29  (0.50) |
| 5 | 11.06  (0.15) | 10.03  (0.24) | 5.13  (0.39) | 8.08  (0.38) | 10.57  (0.29) | 0.39  (0.71) |
| 6 | 11.13  (0.16) | -11.01  ().16) | 7.61  (0.59) | 10.43  (0.45) | -11.26  (0.20) | 9.81  (0.55) |
| 7 | -0.47  (0.44) | 10.77  (0.17) | -4.98  (0.30) | 2.12  (0.54) | 9.66  (0.48) | -1.51  (0.45) |
| 8 | 10.90  (0.29) | -10.64  (0.26) | 1.37  (0.48) | 1.57  (0.89) | -2.09  (0.68) | -0.09  (0.81) |
| 9 | 8.25  (0.42) | 11.66  (0.10) | 7.39  (0.54) | -0.74  (1.09) | 11.03  (0.23) | 1.77  (0.90) |
| 10 | 11.35  (0.23) | -10.42  (0.35) | 5.33  (0.67) | 10.79  (0.37) | 10.21  (0.19) | 0.45  (0.34) |

*Note*: DA = dopamine, 5-HT = serotonin, NE = norepinephrine.

#### Supplementary Table 2. Summary statistics of session-level behavior across tasks.

| **Session** | **Task** | **Behavior** | **N** | **Mean (SD)** |
| --- | --- | --- | --- | --- |
| Baseline | RL | p(Optimal) | 10 | 0.71 (0.08) |
|  |  | RT |  | 1.14 (0.77) |
|  | UG | p(Accept) | 10 | 0.81 (0.31) |
|  |  | RT |  | 2.10 (0.85) |
|  |  | Mood |  | 33.69 (21.06) |
| Pre-Stimulation | RL | p(Optimal) | 10 | 0.73 (0.09) |
|  |  | RT |  | 0.99 (0.69) |
|  | UG | p(Accept) | 10 | 0.62 (0.18) |
|  |  | RT |  | 1.93 (0.51) |
|  |  | Mood |  | 33.24 (17.64) |
| Post-Stimulation | RL | p(Optimal) | 10 | 0.72 (0.08) |
|  |  | RT |  | 0.84 (0.62) |
|  | UG | p(Accept) | 10 | 0.61 (0.22) |
|  |  | RT |  | 1.97 (1.18) |
|  |  | Mood |  | 39.19 (17.07) |
| Week 1 | RL | p(Optimal) | 9 | 0.76 (0.04) |
|  |  | RT |  | 0.56 (0.32) |
|  | UG | p(Accept) | 9 | 0.69 (0.15) |
|  |  | RT |  | 1.36 (0.28) |
|  |  | Mood |  | 44.23 (16.60) |
| Month 1 | RL | p(Optimal) | 10 | 0.72 (0.06) |
|  |  | RT |  | 0.46 (0.27) |
|  | UG | p(Accept) | 10 | 0.64 (0.13) |
|  |  | RT |  | 1.31 (0.41) |
|  |  | Mood |  | 46.95 (16.74) |
| Month 2 | RL | p(Optimal) | 9 | 0.75 (0.02) |
|  |  | RT |  | 0.44 (0.26) |
|  | UG | p(Accept) | 9 | 0.66 (0.22) |
|  |  | RT |  | 1.23 (0.32) |
|  |  | Mood |  | 45.60 (17.51) |
| Month 3 | RL | p(Optimal) | 10 | 0.73 (0.08) |
|  |  | RT |  | 0.55 (0.35) |
|  | UG | p(Accept) | 10 | 0.62 (0.22) |
|  |  | RT |  | 1.26 (0.20) |
|  |  | Mood |  | 49.02 (8.64) |
| Month 4 | RL | p(Optimal) | 8 | 0.75 (0.05) |
|  |  | RT |  | 0.59 (0.44) |
|  | UG | p(Accept) | 9 | 0.68 (0.21) |
|  |  | RT |  | 1.35 (0.43) |
|  |  | Mood |  | 48.26 (11.99) |
| Month 5 | RL | p(Optimal) | 8 | 0.69 (0.08) |
|  |  | RT |  | 0.36 (0.26) |
|  | UG | p(Accept) | 8 | 0.58 (0.24) |
|  |  | RT |  | 1.25 (0.24) |
|  |  | Mood |  | 49.50 (8.88) |
| Month 6 | RL | p(Optimal) | 10 | 0.76 (0.06) |
|  |  | RT |  | 0.45 (0.28) |
|  | UG | p(Accept) | 10 | 0.55 (0.19) |
|  |  | RT |  | 1.30 (0.30) |
|  |  | Mood |  | 54.31 (7.49) |

*Note:* RL = reversal learning task, UG = ultimatum game, p(Optimal) = probability of choosing the more rewarding option, p(Accept) = probability of accepting an offer, RT = response time (given in seconds), Mood = average mood rating (0-100 scale).

#### Supplementary Table 3. Neural models of acute stimulation in reversal learning (N-RL-M0).

| **Neurotransmitter** | **Effect** | **Est.** | **SE** | **T** | **P** |
| --- | --- | --- | --- | --- | --- |
| Dopamine | Intercept | -3.24 | 1.35 | -2.40 | 0.043* |
| Dopamine | Stimulation Session (Pre vs. Post) | 6.41 | 2.71 | 2.37 | 0.046* |
| Dopamine | Trial Number (z-scored) | 0.74 | 0.09 | 8.22 | < 0.001*** |
| Dopamine | Stim × Trial | -1.25 | 0.13 | -9.74 | < 0.001*** |
| Serotonin | Intercept | -0.64 | 1.78 | -0.36 | 0.730 |
| Serotonin | Stimulation Session (Pre vs. Post) | 1.31 | 3.55 | 0.37 | 0.722 |
| Serotonin | Trial Number (z-scored) | 0.48 | 0.08 | 6.08 | < 0.001*** |
| Serotonin | Stim × Trial | 0.10 | 0.11 | 0.87 | 0.385 |
| Norepinephrine | Intercept | 0.34 | 1.19 | 0.29 | 0.783 |
| Norepinephrine | Stimulation Session (Pre vs. Post) | -0.57 | 2.39 | -0.24 | 0.819 |
| Norepinephrine | Trial Number (z-scored) | -1.51 | 0.12 | -12.66 | < 0.001*** |
| Norepinephrine | Stim × Trial | 4.06 | 0.17 | 23.96 | < 0.001*** |

*Note*: † = P < 0.10, * = P < 0.05, ** = P < 0.01, *** = P < 0.001.

#### Supplementary Table 4. Neural models of acute stimulation in the ultimatum game (N-UG-M0).

| **Neurotransmitter** | **Effect** | **Est.** | **SE** | **T** | **P** |
| --- | --- | --- | --- | --- | --- |
| Dopamine | Intercept | -1.41 | 0.95 | -1.48 | 0.171 |
| Dopamine | Stimulation Session (Pre vs. Post) | 3.09 | 1.87 | 1.66 | 0.129 |
| Dopamine | Trial Number (z-scored) | 1.68 | 0.17 | 9.82 | < 0.001*** |
| Dopamine | Stim × Trial | -2.78 | 0.24 | -11.43 | < 0.001*** |
| Serotonin | Intercept | -2.82 | 1.08 | -2.62 | 0.025* |
| Serotonin | Stimulation Session (Pre vs. Post) | 5.62 | 2.18 | 2.57 | 0.028* |
| Serotonin | Trial Number (z-scored) | -0.17 | 0.14 | -1.21 | 0.227 |
| Serotonin | Stim × Trial | 0.14 | 0.19 | 0.70 | 0.483 |
| Norepinephrine | Intercept | 0.43 | 1.01 | 0.42 | 0.683 |
| Norepinephrine | Stimulation Session (Pre vs. Post) | -1.05 | 1.95 | -0.54 | 0.601 |
| Norepinephrine | Trial Number (z-scored) | -0.66 | 0.18 | -3.67 | < 0.001*** |
| Norepinephrine | Stim × Trial | 2.51 | 0.25 | 9.92 | < 0.001*** |

*Note*: † = P < 0.10, * = P < 0.05, ** = P < 0.01, *** = P < 0.001.

#### Supplementary Table 5. Neural models of acute stimulation across tasks (N-B-M0).

| **Neurotransmitter** | **Effect** | **Est.** | **SE** | **T** | **P** |
| --- | --- | --- | --- | --- | --- |
| Dopamine | Intercept | -3.03 | 1.04 | -2.93 | 0.015* |
| Dopamine | Stimulation Session (Pre vs. Post) | 5.99 | 2.07 | 2.89 | 0.016* |
| Dopamine | Task (RL vs. UG) | 1.63 | 0.20 | 8.13 | < 0.001*** |
| Dopamine | Trial Number (z-scored) | 0.75 | 0.10 | 7.60 | < 0.001*** |
| Dopamine | Stim × Task | -2.89 | 0.29 | -9.83 | < 0.001*** |
| Dopamine | Stim × Trial | -1.25 | 0.14 | -9.00 | < 0.001*** |
| Dopamine | Task × Trial | 0.94 | 0.19 | 4.92 | < 0.001*** |
| Dopamine | Stim × Task × Trial | -1.54 | 0.27 | -5.69 | < 0.001*** |
| Serotonin | Intercept | -0.54 | 1.29 | -0.42 | 0.683 |
| Serotonin | Stimulation Session (Pre vs. Post) | 1.09 | 2.58 | 0.42 | 0.682 |
| Serotonin | Task (RL vs. UG) | -2.26 | 0.19 | -11.74 | < 0.001*** |
| Serotonin | Trial Number (z-scored) | 0.49 | 0.09 | 5.19 | < 0.001*** |
| Serotonin | Stim × Task | 4.49 | 0.28 | 15.90 | < 0.001*** |
| Serotonin | Stim × Trial | 0.09 | 0.13 | 0.65 | 0.516 |
| Serotonin | Task × Trial | -0.64 | 0.18 | -3.52 | < 0.001*** |
| Serotonin | Stim × Task × Trial | 0.01 | 0.26 | 0.03 | 0.975 |
| Norepinephrine | Intercept | -0.03 | 0.96 | -0.03 | 0.978 |
| Norepinephrine | Stimulation Session (Pre vs. Post) | 0.18 | 1.94 | 0.10 | 0.926 |
| Norepinephrine | Task (RL vs. UG) | 0.44 | 0.25 | 1.79 | 0.073† |
| Norepinephrine | Trial Number (z-scored) | -1.52 | 0.12 | -12.50 | < 0.001*** |
| Norepinephrine | Stim × Task | -1.21 | 0.36 | -3.33 | < 0.001*** |
| Norepinephrine | Stim × Trial | 4.06 | 0.17 | 23.63 | < 0.001*** |
| Norepinephrine | Task × Trial | 0.86 | 0.24 | 3.62 | < 0.001*** |
| Norepinephrine | Stim × Task × Trial | -1.54 | 0.33 | -4.60 | < 0.001*** |

*Note*: † = P < 0.10, * = P < 0.05, ** = P < 0.01, *** = P < 0.001.

#### Supplementary Table 6. Reversal learning task basic effects models (N-RL-M1).

| **Neurotransmitter** | **Effect** | **Est.** | **SE** | **T** | **P** |
| --- | --- | --- | --- | --- | --- |
| Dopamine | Intercept | -0.215 | 0.26 | -0.82 | 0.414 |
| Dopamine | Outcome (Win vs Loss) | 0.178 | 0.13 | 1.34 | 0.180 |
| Dopamine | Stimulation Session (Pre vs. Post) | 3.203 | 1.38 | 2.32 | 0.047* |
| Dopamine | Block (Negative vs Mixed) | 0.365 | 0.47 | 0.77 | 0.439 |
| Dopamine | Block (Positive vs Mixed) | -0.165 | 0.27 | -0.61 | 0.543 |
| Dopamine | Trial Number (z-scored) | -0.020 | 0.19 | -0.10 | 0.917 |
| Dopamine | Stim × Outcome | 0.067 | 0.13 | 0.51 | 0.612 |
| Dopamine | Stim × Negative vs Mixed | -0.353 | 0.47 | -0.75 | 0.455 |
| Dopamine | Stim × Positive vs Mixed | 0.245 | 0.27 | 0.90 | 0.368 |
| Dopamine | Stim × Trial | -0.491 | 0.19 | -2.55 | 0.011* |
| Serotonin | Intercept | -0.009 | 0.23 | -0.04 | 0.971 |
| Serotonin | Outcome (Win vs Loss) | -0.076 | 0.12 | -0.66 | 0.511 |
| Serotonin | Stimulation Session (Pre vs. Post) | 1.100 | 1.79 | 0.62 | 0.555 |
| Serotonin | Block (Negative vs Mixed) | -0.018 | 0.41 | -0.04 | 0.966 |
| Serotonin | Block (Positive vs Mixed) | 0.233 | 0.24 | 0.98 | 0.326 |
| Serotonin | Trial Number (z-scored) | 0.540 | 0.17 | 3.21 | 0.001** |
| Serotonin | Stim × Outcome | -0.203 | 0.12 | -1.75 | 0.081† |
| Serotonin | Stim × Negative vs Mixed | -0.415 | 0.41 | -1.01 | 0.315 |
| Serotonin | Stim × Positive vs Mixed | -0.547 | 0.24 | -2.30 | 0.021* |
| Serotonin | Stim × Trial | 0.206 | 0.17 | 1.22 | 0.221 |
| Norepinephrine | Intercept | -0.332 | 0.35 | -0.95 | 0.340 |
| Norepinephrine | Outcome (Win vs Loss) | -0.096 | 0.17 | -0.55 | 0.582 |
| Norepinephrine | Stimulation Session (Pre vs. Post) | -0.686 | 1.24 | -0.55 | 0.594 |
| Norepinephrine | Block (Negative vs Mixed) | 0.455 | 0.62 | 0.73 | 0.465 |
| Norepinephrine | Block (Positive vs Mixed) | 0.889 | 0.36 | 2.48 | 0.013* |
| Norepinephrine | Trial Number (z-scored) | 0.340 | 0.25 | 1.34 | 0.181 |
| Norepinephrine | Stim × Outcome | -0.061 | 0.18 | -0.35 | 0.730 |
| Norepinephrine | Stim × Negative vs Mixed | 1.040 | 0.62 | 1.67 | 0.095† |
| Norepinephrine | Stim × Positive vs Mixed | 0.269 | 0.36 | 0.75 | 0.452 |
| Norepinephrine | Stim × Trial | 1.628 | 0.25 | 6.41 | < 0.001*** |

*Note*: † = P < 0.10, * = P < 0.05, ** = P < 0.01, *** = P < 0.001.

#### Supplementary Table 7. Reversal learning task learning dynamics models (N-RL-M2).

| **Neurotransmitter** | **Effect** | **Est.** | **SE** | **T** | **P** |
| --- | --- | --- | --- | --- | --- |
| Dopamine | Intercept | 0.028 | 0.10 | 0.28 | 0.781 |
| Dopamine | Error Trial | -0.210 | 0.14 | -1.46 | 0.145 |
| Dopamine | Stimulation Session (Pre vs. Post) | 3.162 | 1.36 | 2.33 | 0.048* |
| Dopamine | Post-Reversal Period | 0.014 | 0.20 | 0.07 | 0.946 |
| Dopamine | Slow Response Time | -0.012 | 0.13 | -0.09 | 0.926 |
| Dopamine | Trial Number (z-scored) | 0.122 | 0.06 | 1.90 | 0.058† |
| Dopamine | Stim × Error Trial | -0.059 | 0.15 | -0.40 | 0.686 |
| Dopamine | Stim × Post Reversal | 0.389 | 0.20 | 1.93 | 0.054† |
| Dopamine | Stim × Slow RT | 0.029 | 0.13 | 0.23 | 0.819 |
| Dopamine | Stim × Trial | -0.629 | 0.06 | -9.76 | < 0.001*** |
| Serotonin | Intercept | 0.111 | 0.09 | 1.24 | 0.215 |
| Serotonin | Error Trial | 0.037 | 0.13 | 0.29 | 0.771 |
| Serotonin | Stimulation Session (Pre vs. Post) | 0.532 | 1.78 | 0.30 | 0.772 |
| Serotonin | Post-Reversal Period | -0.146 | 0.15 | -0.97 | 0.330 |
| Serotonin | Slow Response Time | -0.162 | 0.12 | -1.39 | 0.164 |
| Serotonin | Trial Number (z-scored) | 0.535 | 0.06 | 9.51 | < 0.001*** |
| Serotonin | Stim × Error Trial | 0.262 | 0.13 | 2.05 | 0.041* |
| Serotonin | Stim × Post Reversal | -0.103 | 0.15 | -0.69 | 0.492 |
| Serotonin | Stim × Slow RT | 0.134 | 0.11 | 1.19 | 0.233 |
| Serotonin | Stim × Trial | 0.046 | 0.06 | 0.82 | 0.413 |
| Norepinephrine | Intercept | -0.018 | 0.13 | -0.13 | 0.893 |
| Norepinephrine | Error Trial | 0.039 | 0.19 | 0.20 | 0.838 |
| Norepinephrine | Stimulation Session (Pre vs. Post) | -0.143 | 1.20 | -0.12 | 0.907 |
| Norepinephrine | Post-Reversal Period | -0.273 | 0.27 | -1.03 | 0.305 |
| Norepinephrine | Slow Response Time | 0.196 | 0.17 | 1.12 | 0.264 |
| Norepinephrine | Trial Number (z-scored) | 0.518 | 0.08 | 6.09 | < 0.001*** |
| Norepinephrine | Stim × Error Trial | 0.108 | 0.19 | 0.56 | 0.575 |
| Norepinephrine | Stim × Post Reversal | -0.389 | 0.27 | -1.46 | 0.144 |
| Norepinephrine | Stim × Slow RT | -0.257 | 0.17 | -1.51 | 0.131 |
| Norepinephrine | Stim × Trial | 2.029 | 0.08 | 23.88 | < 0.001*** |

*Note*: † = P < 0.10, * = P < 0.05, ** = P < 0.01, *** = P < 0.001.

#### Supplementary Table 8. Reversal learning task reward interaction models (N-RL-M3).

| **Neurotransmitter** | **Effect** | **Est.** | **SE** | **T** | **P** |
| --- | --- | --- | --- | --- | --- |
| Dopamine | Intercept | -0.168 | 0.17 | -0.99 | 0.322 |
| Dopamine | Current Reward (-10 vs 0) | 0.404 | 0.28 | 1.44 | 0.150 |
| Dopamine | Current Reward (10 vs 0) | -0.096 | 0.29 | -0.33 | 0.741 |
| Dopamine | Stimulation Session (Pre vs. Post) | 3.173 | 1.36 | 2.34 | 0.047* |
| Dopamine | Previous Reward (-10 vs 0) | 0.366 | 0.28 | 1.29 | 0.196 |
| Dopamine | Previous Reward: (10 vs 0) | -0.243 | 0.30 | -0.82 | 0.411 |
| Dopamine | Trial Number (z-scored) | 0.137 | 0.10 | 1.35 | 0.178 |
| Dopamine | Stim × Rew -10 | -0.607 | 0.28 | -2.16 | 0.031* |
| Dopamine | Stim × Rew 10 | 0.423 | 0.29 | 1.46 | 0.146 |
| Dopamine | Stim × Prev Rew -10 | -0.468 | 0.28 | -1.65 | 0.099† |
| Dopamine | Stim × Prev Rew 10 | 0.235 | 0.30 | 0.80 | 0.426 |
| Dopamine | Both Rewards -10 | -0.609 | 0.44 | -1.39 | 0.164 |
| Dopamine | Current +10, Previous -10 | -0.153 | 0.46 | -0.33 | 0.740 |
| Dopamine | Current -10, Previous +10 | -0.142 | 0.47 | -0.30 | 0.761 |
| Dopamine | Both Rewards +10 | 0.553 | 0.39 | 1.42 | 0.156 |
| Dopamine | Stim × Trial | -0.458 | 0.10 | -4.50 | < 0.001*** |
| Dopamine | Stim × Rew -10 × Prev Rew -10 | 0.865 | 0.44 | 1.98 | 0.048* |
| Dopamine | Stim × Rew 10 × Prev Rew -10 | 0.476 | 0.46 | 1.03 | 0.302 |
| Dopamine | Stim × Rew -10 × Prev Rew 10 | 0.897 | 0.47 | 1.92 | 0.056† |
| Dopamine | Stim × Rew 10 × Prev Rew 10 | -0.591 | 0.39 | -1.51 | 0.130 |
| Serotonin | Intercept | 0.034 | 0.15 | 0.24 | 0.813 |
| Serotonin | Current Reward (-10 vs 0) | -0.148 | 0.16 | -0.91 | 0.363 |
| Serotonin | Current Reward (10 vs 0) | 0.039 | 0.18 | 0.22 | 0.825 |
| Serotonin | Stimulation Session (Pre vs. Post) | 0.795 | 1.78 | 0.45 | 0.667 |
| Serotonin | Previous Reward (-10 vs 0) | -0.229 | 0.20 | -1.14 | 0.255 |
| Serotonin | Previous Reward: (10 vs 0) | 0.236 | 0.23 | 1.01 | 0.310 |
| Serotonin | Trial Number (z-scored) | 0.574 | 0.08 | 6.82 | < 0.001*** |
| Serotonin | Stim × Rew -10 | 0.006 | 0.16 | 0.04 | 0.969 |
| Serotonin | Stim × Rew 10 | -0.271 | 0.18 | -1.54 | 0.125 |
| Serotonin | Stim × Prev Rew -10 | 0.157 | 0.20 | 0.78 | 0.437 |
| Serotonin | Stim × Prev Rew 10 | -0.108 | 0.23 | -0.46 | 0.642 |
| Serotonin | Both Rewards -10 | -0.016 | 0.02 | -0.77 | 0.442 |
| Serotonin | Current +10, Previous -10 | -0.026 | 0.02 | -1.30 | 0.195 |
| Serotonin | Current -10, Previous +10 | -0.069 | 0.08 | -0.82 | 0.411 |
| Serotonin | Both Rewards +10 | -0.011 | 0.02 | -0.55 | 0.585 |
| Serotonin | Stim × Trial | -0.011 | 0.02 | -0.54 | 0.590 |
| Norepinephrine | Intercept | 0.202 | 0.23 | 0.90 | 0.369 |
| Norepinephrine | Current Reward (-10 vs 0) | -0.116 | 0.37 | -0.31 | 0.756 |
| Norepinephrine | Current Reward (10 vs 0) | -0.137 | 0.39 | -0.35 | 0.723 |
| Norepinephrine | Stimulation Session (Pre vs. Post) | -0.407 | 1.22 | -0.33 | 0.747 |
| Norepinephrine | Previous Reward (-10 vs 0) | -0.537 | 0.38 | -1.43 | 0.153 |
| Norepinephrine | Previous Reward: (10 vs 0) | 0.027 | 0.39 | 0.07 | 0.944 |
| Norepinephrine | Trial Number (z-scored) | 0.442 | 0.14 | 3.27 | 0.001** |
| Norepinephrine | Stim × Rew -10 | 0.575 | 0.37 | 1.54 | 0.123 |
| Norepinephrine | Stim × Rew 10 | -0.192 | 0.39 | -0.50 | 0.619 |
| Norepinephrine | Stim × Prev Rew -10 | 0.641 | 0.38 | 1.70 | 0.089† |
| Norepinephrine | Stim × Prev Rew 10 | -0.088 | 0.39 | -0.22 | 0.823 |
| Norepinephrine | Both Rewards -10 | 0.429 | 0.58 | 0.74 | 0.461 |
| Norepinephrine | Current +10, Previous -10 | -0.047 | 0.61 | -0.08 | 0.939 |
| Norepinephrine | Current -10, Previous +10 | -0.557 | 0.62 | -0.89 | 0.371 |
| Norepinephrine | Both Rewards +10 | 0.216 | 0.52 | 0.42 | 0.677 |
| Norepinephrine | Stim × Trial | 1.920 | 0.14 | 14.19 | < 0.001*** |
| Norepinephrine | Stim × Rew -10 × Prev Rew -10 | -0.848 | 0.58 | -1.46 | 0.145 |
| Norepinephrine | Stim × Rew 10 × Prev Rew -10 | -0.575 | 0.61 | -0.94 | 0.348 |
| Norepinephrine | Stim × Rew -10 × Prev Rew 10 | -0.913 | 0.62 | -1.47 | 0.142 |
| Norepinephrine | Stim × Rew 10 × Prev Rew 10 | 0.438 | 0.52 | 0.84 | 0.398 |

*Note*: † = P < 0.10, * = P < 0.05, ** = P < 0.01, *** = P < 0.001.

#### Supplementary Table 9. Reversal learning task prediction error models (N-RL-M4).

| **Neurotransmitter** | **Effect** | **Est.** | **SE** | **T** | **P** |
| --- | --- | --- | --- | --- | --- |
| Dopamine | Intercept | -0.040 | 0.08 | -0.47 | 0.638 |
| Dopamine | Negative Prediction Error | -0.068 | 0.16 | -0.41 | 0.678 |
| Dopamine | Positive Prediction Error | -0.012 | 0.16 | -0.07 | 0.942 |
| Dopamine | Stimulation Session (Pre vs. Post) | 3.162 | 1.35 | 2.34 | 0.047* |
| Dopamine | Trial Number (z-scored) | 0.162 | 0.06 | 2.54 | 0.011* |
| Dopamine | Stim × Negative PE | 0.053 | 0.16 | 0.33 | 0.744 |
| Dopamine | Stim × Positive PE | 0.147 | 0.16 | 0.91 | 0.364 |
| Dopamine | Stim × Trial Number | -0.638 | 0.06 | -10.00 | < 0.001*** |
| Serotonin | Intercept | -0.039 | 0.07 | -0.52 | 0.602 |
| Serotonin | Negative Prediction Error | 0.138 | 0.15 | 0.95 | 0.343 |
| Serotonin | Positive Prediction Error | 0.114 | 0.14 | 0.79 | 0.430 |
| Serotonin | Stimulation Session (Pre vs. Post) | 0.655 | 1.78 | 0.37 | 0.722 |
| Serotonin | Trial Number (z-scored) | 0.530 | 0.06 | 9.32 | < 0.001*** |
| Serotonin | Stim × Negative PE | -0.021 | 0.15 | -0.15 | 0.884 |
| Serotonin | Stim × Positive PE | 0.032 | 0.14 | 0.22 | 0.826 |
| Serotonin | Stim × Trial Number | 0.052 | 0.06 | 0.92 | 0.357 |
| Norepinephrine | Intercept | 0.209 | 0.11 | 1.87 | 0.061† |
| Norepinephrine | Negative Prediction Error | -0.243 | 0.22 | -1.12 | 0.261 |
| Norepinephrine | Positive Prediction Error | -0.460 | 0.21 | -2.15 | 0.032* |
| Norepinephrine | Stimulation Session (Pre vs. Post) | -0.263 | 1.21 | -0.22 | 0.833 |
| Norepinephrine | Trial Number (z-scored) | 0.467 | 0.08 | 5.52 | < 0.001*** |
| Norepinephrine | Stim × Negative PE | -0.105 | 0.22 | -0.49 | 0.627 |
| Norepinephrine | Stim × Positive PE | -0.034 | 0.21 | -0.16 | 0.873 |
| Norepinephrine | Stim × Trial Number | 2.043 | 0.08 | 24.14 | < 0.001*** |

*Note*: † = P < 0.10, * = P < 0.05, ** = P < 0.01, *** = P < 0.001.

#### Supplementary Table 10. Ultimatum game task basic effects models (N-UG-M1).

| **Neurotransmitter** | **Effect** | **Est.** | **SE** | **T** | **P** |
| --- | --- | --- | --- | --- | --- |
| Dopamine | Intercept | -1.049 | 0.98 | -1.08 | 0.305 |
| Dopamine | Offer (Low vs Middle) | -1.154 | 0.41 | -2.80 | 0.005** |
| Dopamine | Offer (High vs Middle) | -0.184 | 0.41 | -0.45 | 0.656 |
| Dopamine | Stimulation Session (Pre vs. Post) | 2.950 | 1.89 | 1.56 | 0.148 |
| Dopamine | Trial Number (z-scored) | 1.666 | 0.17 | 9.74 | < 0.001*** |
| Dopamine | Stim × Low Offer | 0.963 | 0.58 | 1.65 | 0.099† |
| Dopamine | Stim × Stimulation | -0.435 | 0.59 | -0.74 | 0.460 |
| Dopamine | Stim × Trial Number | -2.734 | 0.24 | -11.28 | < 0.001*** |
| Serotonin | Intercept | -2.937 | 1.09 | -2.70 | 0.021* |
| Serotonin | Offer (Low vs Middle) | 0.346 | 0.33 | 1.04 | 0.300 |
| Serotonin | Offer (High vs Middle) | 0.080 | 0.34 | 0.24 | 0.810 |
| Serotonin | Stimulation Session (Pre vs. Post) | 5.767 | 2.19 | 2.63 | 0.025* |
| Serotonin | Trial Number (z-scored) | -0.158 | 0.14 | -1.14 | 0.255 |
| Serotonin | Stim × Low Offer | -0.455 | 0.47 | -0.96 | 0.335 |
| Serotonin | Stim × Stimulation | -0.110 | 0.48 | -0.23 | 0.818 |
| Serotonin | Stim × Trial Number | 0.136 | 0.20 | 0.69 | 0.488 |
| Norepinephrine | Intercept | 0.264 | 1.03 | 0.26 | 0.803 |
| Norepinephrine | Offer (Low vs Middle) | 0.549 | 0.43 | 1.27 | 0.204 |
| Norepinephrine | Offer (High vs Middle) | 0.054 | 0.43 | 0.12 | 0.902 |
| Norepinephrine | Stimulation Session (Pre vs. Post) | -0.988 | 1.98 | -0.50 | 0.627 |
| Norepinephrine | Trial Number (z-scored) | -0.650 | 0.18 | -3.62 | < 0.001*** |
| Norepinephrine | Stim × Low Offer | -0.646 | 0.61 | -1.06 | 0.292 |
| Norepinephrine | Stim × Stimulation | 0.409 | 0.62 | 0.66 | 0.508 |
| Norepinephrine | Stim × Trial Number | 2.490 | 0.25 | 9.78 | < 0.001*** |

*Note*: † = P < 0.10, * = P < 0.05, ** = P < 0.01, *** = P < 0.001

#### Supplementary Table 11. Ultimatum game task context effects models (N-UG-M2).

| **Neurotransmitter** | **Effect** | **Est.** | **SE** | **T** | **P** |
| --- | --- | --- | --- | --- | --- |
| Dopamine | Intercept | -1.725 | 1.01 | -1.71 | 0.113 |
| Dopamine | Choice (Accept vs Reject) | 0.321 | 0.37 | 0.87 | 0.386 |
| Dopamine | Stimulation Session (Pre vs. Post) | 3.442 | 1.93 | 1.79 | 0.101 |
| Dopamine | Slow Response Time | 0.217 | 0.36 | 0.60 | 0.546 |
| Dopamine | Trial Number (z-scored) | 1.679 | 0.17 | 9.75 | < 0.001*** |
| Dopamine | Stim × Accept | -0.251 | 0.55 | -0.46 | 0.646 |
| Dopamine | Stim × Slow RT | -0.381 | 0.50 | -0.76 | 0.446 |
| Dopamine | Stim × Trial Number | -2.773 | 0.24 | -11.38 | < 0.001*** |
| Serotonin | Intercept | -2.152 | 1.12 | -1.92 | 0.081† |
| Serotonin | Choice (Accept vs Reject) | -0.363 | 0.29 | -1.24 | 0.217 |
| Serotonin | Stimulation Session (Pre vs. Post) | 4.614 | 2.22 | 2.08 | 0.063† |
| Serotonin | Slow Response Time | -0.816 | 0.29 | -2.86 | 0.004** |
| Serotonin | Trial Number (z-scored) | -0.164 | 0.14 | -1.19 | 0.233 |
| Serotonin | Stim × Accept | 0.618 | 0.44 | 1.42 | 0.157 |
| Serotonin | Stim × Slow RT | 1.205 | 0.40 | 3.03 | 0.003** |
| Serotonin | Stim × Trial Number | 0.126 | 0.19 | 0.65 | 0.515 |
| Norepinephrine | Intercept | 0.686 | 1.07 | 0.64 | 0.534 |
| Norepinephrine | Choice (Accept vs Reject) | -0.404 | 0.39 | -1.05 | 0.295 |
| Norepinephrine | Stimulation Session (Pre vs. Post) | -1.610 | 2.02 | -0.80 | 0.442 |
| Norepinephrine | Slow Response Time | -0.020 | 0.37 | -0.05 | 0.958 |
| Norepinephrine | Trial Number (z-scored) | -0.648 | 0.18 | -3.61 | < 0.001*** |
| Norepinephrine | Stim × Accept | 0.531 | 0.57 | 0.93 | 0.352 |
| Norepinephrine | Stim × Slow RT | 0.504 | 0.52 | 0.97 | 0.333 |
| Norepinephrine | Stim × Trial Number | 2.494 | 0.25 | 9.82 | < 0.001*** |

*Note*: † = P < 0.10, * = P < 0.05, ** = P < 0.01, *** = P < 0.001

#### Supplementary Table 12. Ultimatum game task offer interactions models (N-UG-M3).

| **Neurotransmitter** | **Effect** | **Est.** | **SE** | **T** | **P** |
| --- | --- | --- | --- | --- | --- |
| Dopamine | Intercept | -0.831 | 1.00 | -0.83 | 0.420 |
| Dopamine | Offer (Low vs Middle) | -1.433 | 0.54 | -2.65 | 0.008** |
| Dopamine | Offer (High vs Middle) | -0.353 | 0.56 | -0.64 | 0.525 |
| Dopamine | Stimulation Session (Post vs Pre) | 2.693 | 1.90 | 1.42 | 0.183 |
| Dopamine | Previous Offer (Low vs Middle) | -0.393 | 0.60 | -0.65 | 0.515 |
| Dopamine | Previous Offer (High vs Middle) | -0.523 | 0.66 | -0.79 | 0.431 |
| Dopamine | Trial Number (z-scored) | 1.667 | 0.18 | 9.40 | < 0.001*** |
| Dopamine | Stim × Low Offer | 1.014 | 0.76 | 1.34 | 0.181 |
| Dopamine | Stim × High Offer | -0.242 | 0.78 | -0.31 | 0.757 |
| Dopamine | Stim × Previous Low Offer | 0.936 | 0.85 | 1.10 | 0.270 |
| Dopamine | Stim × Previous High Offer | -0.117 | 0.96 | -0.12 | 0.904 |
| Dopamine | Low Offer × Previous Low | -0.094 | 0.98 | -0.10 | 0.923 |
| Dopamine | High Offer × Previous Low | 0.093 | 0.97 | 0.10 | 0.924 |
| Dopamine | Low Offer × Previous High | 1.846 | 1.17 | 1.58 | 0.115 |
| Dopamine | High Offer × Previous High | 0.733 | 1.13 | 0.65 | 0.517 |
| Dopamine | Stim × Trial Number | -2.729 | 0.25 | -10.92 | < 0.001*** |
| Dopamine | Stim × Low × PrevLow | 0.326 | 1.40 | 0.23 | 0.815 |
| Dopamine | Stim × High × PrevLow | -0.802 | 1.35 | -0.59 | 0.554 |
| Dopamine | Stim × Low × PrevHigh | -0.649 | 1.64 | -0.40 | 0.692 |
| Dopamine | Stim × High × PrevHigh | 0.355 | 1.65 | 0.21 | 0.830 |
| Serotonin | Intercept | -2.990 | 1.11 | -2.69 | 0.021* |
| Serotonin | Offer (Low vs Middle) | 0.470 | 0.44 | 1.06 | 0.288 |
| Serotonin | Offer (High vs Middle) | -0.085 | 0.45 | -0.19 | 0.852 |
| Serotonin | Stimulation Session (Post vs Pre) | 5.683 | 2.21 | 2.58 | 0.027* |
| Serotonin | Previous Offer (Low vs Middle) | 0.051 | 0.49 | 0.10 | 0.918 |
| Serotonin | Previous Offer (High vs Middle) | 0.016 | 0.54 | 0.03 | 0.976 |
| Serotonin | Trial Number (z-scored) | -0.135 | 0.15 | -0.93 | 0.351 |
| Serotonin | Stim × Low Offer | -0.387 | 0.62 | -0.62 | 0.533 |
| Serotonin | Stim × High Offer | 0.043 | 0.64 | 0.07 | 0.946 |
| Serotonin | Stim × Previous Low Offer | 0.001 | 0.69 | 0.00 | 0.999 |
| Serotonin | Stim × Previous High Offer | 0.406 | 0.79 | 0.51 | 0.607 |
| Serotonin | Low Offer × Previous Low | -0.121 | 0.80 | -0.15 | 0.880 |
| Serotonin | High Offer × Previous Low | 0.847 | 0.79 | 1.07 | 0.285 |
| Serotonin | Low Offer × Previous High | -0.400 | 0.96 | -0.42 | 0.675 |
| Serotonin | High Offer × Previous High | -0.232 | 0.93 | -0.25 | 0.802 |
| Serotonin | Stim × Trial Number | 0.149 | 0.20 | 0.73 | 0.467 |
| Serotonin | Stim × Low × PrevLow | -0.083 | 1.14 | -0.07 | 0.942 |
| Serotonin | Stim × High × PrevLow | -0.505 | 1.11 | -0.46 | 0.649 |
| Serotonin | Stim × Low × PrevHigh | -0.167 | 1.34 | -0.12 | 0.901 |
| Serotonin | Stim × High × PrevHigh | -0.042 | 1.35 | -0.03 | 0.975 |
| Norepinephrine | Intercept | -0.114 | 1.08 | -0.11 | 0.917 |
| Norepinephrine | Offer (Low vs Middle) | 1.032 | 0.56 | 1.84 | 0.067† |
| Norepinephrine | Offer (High vs Middle) | 0.817 | 0.58 | 1.42 | 0.157 |
| Norepinephrine | Stimulation Session (Post vs Pre) | -0.721 | 2.03 | -0.36 | 0.729 |
| Norepinephrine | Previous Offer (Low vs Middle) | 0.480 | 0.63 | 0.77 | 0.444 |
| Norepinephrine | Previous Offer (High vs Middle) | 1.584 | 0.69 | 2.30 | 0.022* |
| Norepinephrine | Trial Number (z-scored) | -0.711 | 0.18 | -3.86 | < 0.001*** |
| Norepinephrine | Stim × Low Offer | -0.921 | 0.79 | -1.17 | 0.242 |
| Norepinephrine | Stim × High Offer | 0.300 | 0.81 | 0.37 | 0.712 |
| Norepinephrine | Stim × Previous Low Offer | -0.570 | 0.88 | -0.65 | 0.517 |
| Norepinephrine | Stim × Previous High Offer | -0.707 | 1.00 | -0.71 | 0.481 |
| Norepinephrine | Low Offer × Previous Low | 0.166 | 1.02 | 0.16 | 0.871 |
| Norepinephrine | High Offer × Previous Low | -1.620 | 1.01 | -1.61 | 0.108 |
| Norepinephrine | Low Offer × Previous High | -3.552 | 1.21 | -2.93 | 0.004** |
| Norepinephrine | High Offer × Previous High | -1.951 | 1.17 | -1.66 | 0.097† |
| Norepinephrine | Stim × Trial Number | 2.524 | 0.26 | 9.73 | < 0.001*** |
| Norepinephrine | Stim × Low × PrevLow | -1.250 | 1.45 | -0.86 | 0.389 |
| Norepinephrine | Stim × High × PrevLow | 0.795 | 1.41 | 0.57 | 0.572 |
| Norepinephrine | Stim × Low × PrevHigh | 3.710 | 1.70 | 2.18 | 0.029* |
| Norepinephrine | Stim × High × PrevHigh | -0.755 | 1.72 | -0.44 | 0.660 |

*Note*: † = P < 0.10, * = P < 0.05, ** = P < 0.01, *** = P < 0.001.

#### Supplementary Table 13. Ultimatum game task offer dynamics models (N-UG-M4).

| **Neurotransmitter** | **Effect** | **Est.** | **SE** | **T** | **P** |
| --- | --- | --- | --- | --- | --- |
| Dopamine | Intercept | -1.396 | 1.07 | -1.30 | 0.211 |
| Dopamine | Offer Change (Worse vs Same) | -0.062 | 0.58 | -0.11 | 0.915 |
| Dopamine | Offer Change (Better vs Same) | 0.017 | 0.58 | 0.03 | 0.976 |
| Dopamine | Stimulation Session (Post vs Pre) | 3.175 | 2.00 | 1.59 | 0.135 |
| Dopamine | Trial Number (z-scored) | 1.703 | 0.18 | 9.59 | < 0.001*** |
| Dopamine | Stim × Worse Offer | -0.230 | 0.85 | -0.27 | 0.786 |
| Dopamine | Stim × Better Offer | 0.054 | 0.85 | 0.06 | 0.949 |
| Dopamine | Stim × Trial Number | -2.791 | 0.25 | -11.10 | < 0.001*** |
| Serotonin | Intercept | -2.598 | 1.15 | -2.26 | 0.042* |
| Serotonin | Offer Change (Worse vs Same) | -0.304 | 0.47 | -0.65 | 0.516 |
| Serotonin | Offer Change (Better vs Same) | -0.226 | 0.47 | -0.48 | 0.633 |
| Serotonin | Stimulation Session (Post vs Pre) | 5.314 | 2.25 | 2.36 | 0.037* |
| Serotonin | Trial Number (z-scored) | -0.145 | 0.14 | -1.01 | 0.313 |
| Serotonin | Stim × Worse Offer | 0.472 | 0.68 | 0.69 | 0.491 |
| Serotonin | Stim × Better Offer | 0.166 | 0.69 | 0.24 | 0.809 |
| Serotonin | Stim × Trial Number | 0.166 | 0.20 | 0.82 | 0.414 |
| Norepinephrine | Intercept | 0.927 | 1.15 | 0.80 | 0.433 |
| Norepinephrine | Offer Change (Worse vs Same) | -0.389 | 0.60 | -0.65 | 0.519 |
| Norepinephrine | Offer Change (Better vs Same) | -0.666 | 0.61 | -1.09 | 0.275 |
| Norepinephrine | Stimulation Session (Post vs Pre) | -2.221 | 2.14 | -1.04 | 0.317 |
| Norepinephrine | Trial Number (z-scored) | -0.735 | 0.19 | -3.97 | < 0.001*** |
| Norepinephrine | Stim × Worse Offer | 1.199 | 0.88 | 1.36 | 0.175 |
| Norepinephrine | Stim × Better Offer | 1.352 | 0.89 | 1.53 | 0.128 |
| Norepinephrine | Stim × Trial Number | 2.568 | 0.26 | 9.79 | < 0.001*** |

*Note*: † = P < 0.10, * = P < 0.05, ** = P < 0.01, *** = P < 0.001.

#### Supplementary Table 14. Ultimatum game choice × session models (Batten-M1).

| **Neurotransmitter** | **Effect** | **Est.** | **SE** | **T** | **P** |
| --- | --- | --- | --- | --- | --- |
| Dopamine | Intercept | -1.73 | 0.98 | -1.76 | 0.105 |
| Dopamine | Stimulation Session (Post vs Pre) | 3.38 | 1.90 | 1.78 | 0.103 |
| Dopamine | Choice (Accept vs Reject) | 0.53 | 0.40 | 1.30 | 0.193 |
| Dopamine | Stim × Choice | -0.47 | 0.59 | -0.79 | 0.428 |
| Serotonin | Intercept | -2.66 | 1.09 | -2.45 | 0.033* |
| Serotonin | Stimulation Session (Post vs Pre) | 5.38 | 2.19 | 2.45 | 0.033* |
| Serotonin | Choice (Accept vs Reject) | -0.26 | 0.29 | -0.90 | 0.368 |
| Serotonin | Stim × Choice | 0.38 | 0.43 | 0.88 | 0.368 |
| Norepinephrine | Intercept | 0.73 | 1.05 | 0.69 | 0.503 |
| Norepinephrine | Stimulation Session (Post vs Pre) | -1.42 | 2.00 | -0.71 | 0.492 |
| Norepinephrine | Choice (Accept vs Reject) | -0.49 | 0.42 | -1.17 | 0.241 |
| Norepinephrine | Stim × Choice | 0.59 | 0.61 | 0.97 | 0.335 |

*Note*: † = P < 0.10, * = P < 0.05, ** = P < 0.01, *** = P < 0.001.

#### Supplementary Table 15. Ultimatum game task offer value models (Batten-M3).

| **Neurotransmitter** | **Effect** | **Est.** | **SE** | **T** | **P** |
| --- | --- | --- | --- | --- | --- |
| Dopamine | Intercept | -0.80 | 0.55 | -1.47 | 0.142 |
| Dopamine | Offer Value | 0.15 | 0.10 | 1.44 | 0.149 |
| Dopamine | Offer Value Difference | -0.09 | 0.07 | -1.17 | 0.241 |
| Serotonin | Intercept | -0.48 | 0.71 | -0.68 | 0.512 |
| Serotonin | Offer Value | -0.01 | 0.11 | -0.05 | 0.962 |
| Serotonin | Offer Value Difference | 0.05 | 0.06 | 0.75 | 0.456 |
| Norepinephrine | Intercept | 0.37 | 0.51 | 0.72 | 0.478 |
| Norepinephrine | Offer Value | -0.04 | 0.10 | -0.37 | 0.709 |
| Norepinephrine | Offer Value Difference | -0.08 | 0.07 | -1.13 | 0.261 |

*Note*: † = P < 0.10, * = P < 0.05, ** = P < 0.01, *** = P < 0.001.

#### Supplementary Table 16. Ultimatum game task offer value × session models (Batten-M3-Stim).

| **Neurotransmitter** | **Effect** | **Est.** | **SE** | **T** | **P** |
| --- | --- | --- | --- | --- | --- |
| Dopamine | Intercept | -1.21 | 0.79 | -1.54 | 0.126 |
| Dopamine | Offer Value | 0.25 | 0.14 | 1.74 | 0.082† |
| Dopamine | Stimulation Session (Post vs Pre) | 0.85 | 1.14 | 0.74 | 0.458 |
| Dopamine | Offer Value Difference | -0.13 | 0.10 | -1.28 | 0.201 |
| Dopamine | Offer Value × Stim | -0.21 | 0.21 | -1.01 | 0.312 |
| Dopamine | Value Difference × Stim | 0.09 | 0.14 | 0.63 | 0.528 |
| Serotonin | Intercept | -0.48 | 0.70 | -0.68 | 0.497 |
| Serotonin | Offer Value | -0.05 | 0.12 | -0.42 | 0.673 |
| Serotonin | Stimulation Session (Post vs Pre) | 0.01 | 0.91 | 0.02 | 0.987 |
| Serotonin | Offer Value Difference | 0.15 | 0.08 | 1.81 | 0.071† |
| Serotonin | Offer Value × Stim | 0.09 | 0.17 | 0.52 | 0.603 |
| Serotonin | Value Difference × Stim | -0.21 | 0.12 | -1.74 | 0.082† |
| Norepinephrine | Intercept | 0.08 | 0.70 | 0.12 | 0.908 |
| Norepinephrine | Offer Value | 0.03 | 0.13 | 0.22 | 0.826 |
| Norepinephrine | Stimulation Session (Post vs Pre) | 0.59 | 1.00 | 0.59 | 0.553 |
| Norepinephrine | Offer Value Difference | -0.14 | 0.09 | -1.52 | 0.130 |
| Norepinephrine | Offer Value × Stim | -0.14 | 0.19 | -0.71 | 0.478 |
| Norepinephrine | Value Difference × Stim | 0.13 | 0.13 | 1.00 | 0.319 |

*Note*: † = P < 0.10, * = P < 0.05, ** = P < 0.01, *** = P < 0.001.

#### Supplementary Table 17. Ultimatum game task modified offer value models (Batten-M3*).

| **Neurotransmitter** | **Effect** | **Est.** | **SE** | **T** | **P** |
| --- | --- | --- | --- | --- | --- |
| Dopamine | Intercept | -0.35 | 0.39 | -0.91 | 0.366 |
| Dopamine | Offer Value | 0.06 | 0.07 | 0.86 | 0.392 |
| Dopamine | Offer Value Difference (Residual) | -0.09 | 0.07 | -1.23 | 0.218 |
| Serotonin | Intercept | -0.72 | 0.67 | -1.07 | 0.308 |
| Serotonin | Offer Value | 0.04 | 0.09 | 0.47 | 0.649 |
| Serotonin | Offer Value Difference (Residual) | 0.04 | 0.06 | 0.70 | 0.484 |
| Norepinephrine | Intercept | 0.77 | 0.36 | 2.14 | 0.036* |
| Norepinephrine | Offer Value | -0.12 | 0.06 | -1.80 | 0.073† |
| Norepinephrine | Offer Value Difference (Residual) | -0.07 | 0.07 | -1.05 | 0.297 |

*Note*: † = P < 0.10, * = P < 0.05, ** = P < 0.01, *** = P < 0.001.

#### Supplementary Table 18. Ultimatum game task modified offer value × session models (Batten-M3*-Stim).

| **Neurotransmitter** | **Effect** | **Est.** | **SE** | **T** | **P** |
| --- | --- | --- | --- | --- | --- |
| Dopamine | Intercept | -0.54 | 0.59 | -0.92 | 0.362 |
| Dopamine | Offer Value | 0.12 | 0.10 | 1.19 | 0.238 |
| Dopamine | Stimulation Session (Post vs Pre) | 0.37 | 0.86 | 0.43 | 0.666 |
| Dopamine | Offer Value Difference (Residual) | -0.14 | 0.10 | -1.40 | 0.162 |
| Dopamine | Offer Value × Stim | -0.11 | 0.14 | -0.81 | 0.419 |
| Dopamine | Residual Difference × Stim | 0.11 | 0.14 | 0.75 | 0.454 |
| Serotonin | Intercept | -1.28 | 0.55 | -2.35 | 0.023* |
| Serotonin | Offer Value | 0.11 | 0.08 | 1.32 | 0.188 |
| Serotonin | Stimulation Session (Post vs Pre) | 1.12 | 0.66 | 1.71 | 0.089† |
| Serotonin | Offer Value Difference (Residual) | 0.15 | 0.08 | 1.76 | 0.079† |
| Serotonin | Offer Value × Stim | -0.13 | 0.12 | -1.11 | 0.266 |
| Serotonin | Residual Difference × Stim | -0.22 | 0.12 | -1.81 | 0.070† |
| Norepinephrine | Intercept | 0.82 | 0.50 | 1.64 | 0.102 |
| Norepinephrine | Offer Value | -0.12 | 0.09 | -1.30 | 0.195 |
| Norepinephrine | Stimulation Session (Post vs Pre) | -0.11 | 0.72 | -0.15 | 0.883 |
| Norepinephrine | Offer Value Difference (Residual) | -0.14 | 0.09 | -1.46 | 0.145 |
| Norepinephrine | Offer Value × Stim | <0.01 | 0.13 | 0.03 | 0.975 |
| Norepinephrine | Residual Difference × Stim | 0.13 | 0.13 | 1.00 | 0.317 |

*Note*: † = P < 0.10, * = P < 0.05, ** = P < 0.01, *** = P < 0.001.

#### Supplementary Table 19. Ultimatum game task unsigned offer value models (Batten-M4).

| **Neurotransmitter** | **Effect** | **Est.** | **SE** | **T** | **P** |
| --- | --- | --- | --- | --- | --- |
| Dopamine | Intercept | -0.42 | 0.68 | -0.61 | 0.549 |
| Dopamine | Offer Value | 0.14 | 0.11 | 1.35 | 0.181 |
| Dopamine | Offer Value Difference (Signed) | -0.09 | 0.08 | -1.09 | 0.285 |
| Dopamine | Offer Value Difference (Unsigned) | -1.12 | 0.12 | -1.00 | 0.346 |
| Serotonin | Intercept | -0.77 | 0.73 | -1.05 | 0.317 |
| Serotonin | Offer Value | -0.01 | 0.11 | -0.07 | 0.944 |
| Serotonin | Offer Value Difference (Signed) | 0.04 | 0.06 | 0.73 | 0.467 |
| Serotonin | Offer Value Difference (Unsigned) | 0.11 | 0.07 | 1.56 | 0.145 |
| Norepinephrine | Intercept | 0.39 | 0.54 | 0.72 | 0.472 |
| Norepinephrine | Offer Value | -0.04 | 0.10 | -0.37 | 0.716 |
| Norepinephrine | Offer Value Difference (Signed) | -0.07 | 0.07 | -1.07 | 0.290 |
| Norepinephrine | Offer Value Difference (Unsigned) | -0.01 | 0.09 | -0.10 | 0.923 |

*Note*: † = P < 0.10, * = P < 0.05, ** = P < 0.01, *** = P < 0.001.

#### Supplementary Table 20. Ultimatum game task unsigned offer value × session models (Batten-M4-Stim).

| **Neurotransmitter** | **Effect** | **Est.** | **SE** | **T** | **P** |
| --- | --- | --- | --- | --- | --- |
| Dopamine | Intercept | -0.57 | 0.85 | -0.68 | 0.500 |
| Dopamine | Stimulation Session (Post vs Pre) | 0..37 | 1.22 | 0.30 | 0.764 |
| Dopamine | Offer Value | 0.26 | 0.14 | 1.78 | 0.076† |
| Dopamine | Offer Value Difference (Signed) | -0.12 | 0.10 | -1.22 | 0.223 |
| Dopamine | Offer Value Difference (Unsigned) | -0.25 | 0.12 | -2.06 | 0.040* |
| Dopamine | Stim × Value | -0.21 | 0.21 | -1.03 | 0.304 |
| Dopamine | Stim × Signed Value Difference | 0.08 | 0.14 | 0.59 | 0.553 |
| Dopamine | Stim × Unsigned Value Difference | 0.18 | 0.17 | 1.09 | 0.277 |
| Serotonin | Intercept | -0.94 | 0.75 | -1.25 | 0.213 |
| Serotonin | Stimulation Session (Post vs Pre) | 0.35 | 0.98 | 0.36 | 0.722 |
| Serotonin | Offer Value | -0.05 | 0.12 | -0.45 | 0.656 |
| Serotonin | Offer Value Difference (Signed) | 0.15 | 0.08 | 1.76 | 0.079† |
| Serotonin | Offer Value Difference (Unsigned) | 0.18 | 0.10 | 1.75 | 0.081† |
| Serotonin | Stim × Value | 0.09 | 0.17 | 0.53 | 0.596 |
| Serotonin | Stim × Signed Value Difference | -0.20 | 0.12 | -1.71 | 0.088† |
| Serotonin | Stim × Unsigned Value Difference | -0.13 | 0.14 | -0.89 | 0.373 |
| Norepinephrine | Intercept | -0.11 | 0.76 | -0.14 | 0.886 |
| Norepinephrine | Stimulation Session (Post vs Pre) | 1.00 | 1.08 | 0.93 | 0.354 |
| Norepinephrine | Offer Value | 0.03 | 0.13 | 0.21 | 0.833 |
| Norepinephrine | Offer Value Difference (Signed) | -0.14 | 0.09 | -1.54 | 0.125 |
| Norepinephrine | Offer Value Difference (Unsigned) | 0.07 | 0.11 | 0.66 | 0.511 |
| Norepinephrine | Stim × Value | -0.13 | 0.19 | -0.69 | 0.489 |
| Norepinephrine | Stim × Signed Value Difference | 0.13 | 0.13 | 1.01 | 0.311 |
| Norepinephrine | Stim × Unsigned Value Difference | 0.16 | 0.16 | -1.01 | 0.312 |

*Note*: † = P < 0.10, * = P < 0.05, ** = P < 0.01, *** = P < 0.001.

#### Supplementary Table 21. Neurochemical predictions of changes in symptom severity scores.

| **Model** | **AIC** | **BIC** | $\boldsymbol{R}^{\boldsymbol{2}}$ | $\boldsymbol{R}_{\boldsymbol{Adj}}^{\boldsymbol{2}}$ |
| --- | --- | --- | --- | --- |
| ΔDA | 71.09 | 72.00 | 0.03 | -0.09 |
| Δ5-HT | 69.32 | 70.23 | 0.19 | 0.09 |
| ΔNE | 71.32 | 72.23 | 0.01 | -0.12 |
| ΔDA+Δ5-HT | 69.61 | 70.82 | 0.32 | 0.12 |
| ΔNE+Δ5-HT | 70.81 | 72.02 | 0.23 | 0.01 |
| ΔDA+ΔNE | 71.84 | 73.05 | 0.14 | -0.10 |
| ΔDA+Δ5-HT+ΔNE | 71.52 | 73.04 | 0.32 | -0.02 |
| **ΔDA×Δ5-HT** | **55.48** | **56.99** | **0.86** | **0.80** |
| ΔNE×Δ5-HT | 58.99 | 60.50 | 0.81 | 0.71 |
| ΔDA×ΔNE | 62.28 | 63.79 | 0.73 | 0.60 |
| ΔDA×Δ5-HT×ΔNE | 62.15 | 64.87 | 0.88 | 0.46 |

*Note:* DA = dopamine, 5-HT = serotonin, NE = norepinephrine.

#### Supplementary Table 22. Summary of study participants’ stimulation and recording sites in MNI space.

| participant | stimulation site (left) | | | recording site (right) | | |
| --- | --- | --- | --- | --- | --- | --- |
|  | x | y | z | x | y | z |
| 1 | -8.65 | 23.29 | -5.83 | 3.36 | 22.34 | -13.57 |
| 2 | -7.38 | 22.45 | -5.73 | 3.63 | 22.47 | -13.23 |
| 3 | -12.72 | 28.87 | -6.37 | 3.37 | 26.37 | -11.59 |
| 4 | -9.17 | 24.41 | -7.01 | 3.59 | 23.01 | -12.42 |
| 5 | -9.35 | 23.57 | -6.96 | 3.14 | 22.49 | -12.37 |
| 6 | -8.66 | 27.52 | -1.27 | 6.98 | 25.23 | -8.40 |
| 7 | -7.06 | 26.05 | -3.03 | 4.39 | 27.28 | -10.29 |
| 8 | -3.77 | 24.71 | -6.21 | 5.04 | 27.58 | -11.54 |
| 9 | -8.01 | 25.90 | 0.73 | 4.26 | 25.52 | -5.43 |
| 10 | -8.18 | 22.84 | -3.01 | 3.18 | 22.90 | -11.86 |

*Note:* Stimulation site is reported from the left SCC contact (third from the bottom). Recording site is reported from the right SCC contact (lowest).

#### Supplementary Table 23. Statistical tests for stimulation-induced neurotransmitter changes across quantification methods

| Task | Neurotransmitter | Metric | F | P |
| --- | --- | --- | --- | --- |
| Reversal Learning | Dopamine | Sum (500ms) | 24.20 | < 0.001*** |
|  |  | Mean (500ms) | 24.20 | < 0.001*** |
|  |  | Peak (500ms) | 26.66 | < 0.001*** |
|  |  | Relative (500ms) | 1.06 | 0.309 |
|  |  | Sum (1000ms) | 24.10 | < 0.001*** |
|  |  | Mean (1000ms) | 24.10 | < 0.001*** |
|  |  | Peak (1000ms) | 27.41 | < 0.001*** |
|  |  | Relative (1000ms) | 1.31 | 0.258 |
|  | Serotonin | Sum (500ms) | 0.67 | 0.418 |
|  |  | Mean (500ms) | 0.67 | 0.418 |
|  |  | Peak (500ms) | 0.67 | 0.417 |
|  |  | Relative (500ms) | 0.37 | 0.544 |
|  |  | Sum (1000ms) | 0.68 | 0.413 |
|  |  | Mean (1000ms) | 0.68 | 0.413 |
|  |  | Peak (1000ms) | 0.67 | 0.417 |
|  |  | Relative (1000ms) | 0.41 | 0.527 |
|  | Norepinephrine | Sum (500ms) | 0.23 | 0.637 |
|  |  | Mean (500ms) | 0.23 | 0.637 |
|  |  | Peak (500ms) | 0.05 | 0.822 |
|  |  | Relative (500ms) | 0.27 | 0.605 |
|  |  | Sum (1000ms) | 0.24 | 0.629 |
|  |  | Mean (1000ms) | 0.24 | 0.629 |
|  |  | Peak (1000ms) | 0.10 | 0.756 |
|  |  | Relative (1000ms) | 0.14 | 0.708 |
| Ultimatum Game | Dopamine | Sum (500ms) | 13.52 | < 0.001*** |
|  |  | Mean (500ms) | 13.52 | < 0.001*** |
|  |  | Peak (500ms) | 7.74 | 0.007** |
|  |  | Relative (500ms) | 0.07 | 0.786 |
|  |  | Sum (1000ms) | 13.58 | < 0.001*** |
|  |  | Mean (1000ms) | 13.58 | < 0.001*** |
|  |  | Peak (1000ms) | 6.86 | 0.011* |
|  |  | Relative (1000ms) | 0.13 | 0.725 |
|  | Serotonin | Sum (500ms) | 34.58 | < 0.001*** |
|  |  | Mean (500ms) | 34.58 | < 0.001*** |
|  |  | Peak (500ms) | 19.20 | < 0.001*** |
|  |  | Relative (500ms) | 1.66 | 0.203 |
|  |  | Sum (1000ms) | 35.82 | < 0.001*** |
|  |  | Mean (1000ms) | 35.82 | < 0.001*** |
|  |  | Peak (1000ms) | 15.99 | < 0.001*** |
|  |  | Relative (1000ms) | 1.84 | 0.180 |
|  | Norepinephrine | Sum (500ms) | 1.27 | 0.264 |
|  |  | Mean (500ms) | 1.27 | 0.264 |
|  |  | Peak (500ms) | 1.12 | 0.294 |
|  |  | Relative (500ms) | 0.06 | 0.815 |
|  |  | Sum (1000ms) | 1.44 | 0.235 |
|  |  | Mean (1000ms) | 1.44 | 0.235 |
|  |  | Peak (1000ms) | 2.47 | 0.122 |
|  |  | Relative (1000ms) | 0.20 | 0.653 |

*Note:* Two-way repeated-measures ANOVAs testing stimulation effect (Pre vs. Post). RL task includes block type (Reward/Mixed/Punishment); UG task includes offer amount bins. † = P < 0.10, * = P < 0.05, ** = P < 0.01, *** = P < 0.001. df1 = 1 for all stimulation effects. Residual df: RL = 42, UG = 54.

#### Supplementary Table 24. Statistical tests for within-session performance change

| **Task** | **Metric** | **Session** | **Trial Slope (β)** | **SE** | **T/Z** | **P** |
| --- | --- | --- | --- | --- | --- | --- |
| RL | Response time | Pre-Stimulation | -0.061 | 0.034 | -1.80 | 0.073† |
|  |  | Post-Stimulation | 0.077 | 0.034 | 2.37 | 0.023* |
|  | Choice quality | Pre-Stimulation | -0.166 | 0.078 | -2.13 | 0.034* |
|  |  | Post-Stimulation | -0.032 | 0.079 | -0.41 | 0.686 |
| UG | Response time | Pre-Stimulation | -0.018 | 0.024 | -0.77 | 0.442 |
|  |  | Post-Stimulation | 0.026 | 0.024 | 1.10 | 0.274 |
|  | Choice quality | Pre-Stimulation | 0.145 | 0.126 | 1.16 | 0.247 |
|  |  | Post-Stimulation | 0.049 | 0.130 | 0.38 | 0.703 |

*Note:* Trial slopes indicate change per standard deviation increase in trial number. Negative slopes for choice indicate performance decline; positive slopes for RT indicate slowing. Choice quality = optimal choices (RL) or acceptance rate (UG). † = P < 0.10, * = P < 0.05, ** = P < 0.01, *** = P < 0.001.

#### Supplementary Table 25. Pearson correlations between baseline task performance and symptom severity

| **Task** | **Metric** | **r** | **T** | **P** |
| --- | --- | --- | --- | --- |
| RL | Response time | 0.026 | 0.07 | 0.944 |
|  | Choice quality | -0.724 | -2.98 | 0.018* |
| UG | Response time | -0.075 | -0.21 | 0.837 |
|  | Choice quality | 0.151 | 0.43 | 0.676 |
|  | Mood | -0.297 | -0.88 | 0.405 |

*Note:* Trial slopes indicate change per standard deviation increase in trial number. Negative slopes for choice indicate performance decline; positive slopes for RT indicate slowing. Choice quality = optimal choices (RL) or acceptance rate (UG). † = P < 0.10, * = P < 0.05, ** = P < 0.01, *** = P < 0.001.
